# *CRISPRcleanR*^*WebApp*^: an interactive web application for processing genome-wide pooled CRISPR-Cas9 viability screens

**DOI:** 10.1101/2022.03.11.483924

**Authors:** Alessandro Vinceti, Riccardo Roberto de Lucia, Paolo Cremaschi, Umberto Perron, Emre Karacok, Luca Mauri, Carlos Fernandez, Krzysztof Henryk Kluczynski, Daniel Stephen Anderson, Francesco Iorio

## Abstract

A limitation of pooled CRISPR-Cas9 viability screens is the high false-positive rate in detecting *essential genes* arising from copy number-amplified (CNA) regions of the genome. To solve this issue, we developed *CRISPRcleanR*: a computational method implemented as R/python package and in a dockerized version. CRISPRcleanR detects and corrects biased responses to CRISPR-Cas9 targeting in an unsupervised fashion, accurately reducing false-positive signals, while maintaining sensitivity in identifying relevant genetic dependencies. Here, we present *CRISPRcleanR*^*WebApp*^, a web-based application enabling access to CRISPRcleanR through an intuitive graphical web-interface. CRISPRcleanR^WebApp^ removes the complexity of low-level R/python-language user interactions; it provides a user-friendly access to a complete analytical pipeline, not requiring any data pre-processing, and generating gene-level summaries of essentiality with associated statistical scores; it offers a range of interactively explorable plots, while supporting a wider range of CRISPR guide RNAs’ libraries with respect to the original package. CRISPRcleanR^WebApp^ is freely available at: https://crisprcleanr-webapp.fht.org/.

**Highlights:**

- CRISPR-Cas9 screens are widely used for the identification of cancer dependencies
- In such screens, false-positives arise from targeting copy number amplified genes
- CRISPRcleanR corrects this bias in an unsupervised fashion
- CRISPRcleanR^WebApp^ is a web user-friendly front-end for CRISPRcleanR

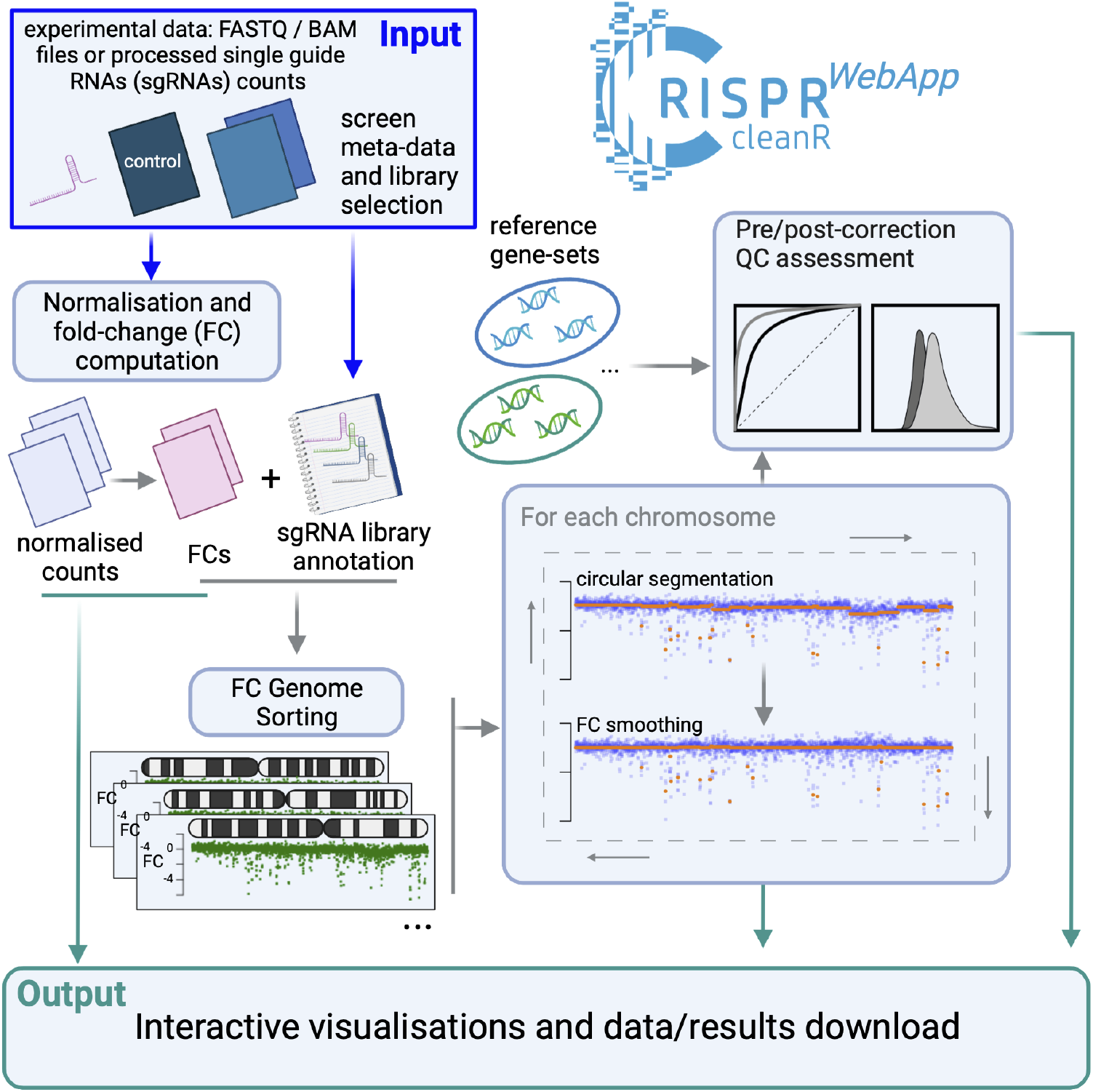

## Introduction

The last decade has seen the advent of genome editing methods based on the Clustered Regularly Interspaced Short Palindromic Repeats (CRISPR) system, which has revolutionised the way molecular biology is investigated and new therapeutic targets are discovered and prioritised (Behan et al., 2019; Cong et al., 2013; Jinek et al., 2012; Mali et al., 2013; Shalem et al., 2014). Particularly, in the last few years, the CRISPR-Cas9 system has been increasingly employed in functional genetic recessive screens, and iteratively optimised reaching unprecedented levels of efficacy, specificity and scalability, thus becoming the state-of-the-art tool for gene inactivation experiments (Barrangou et al., 2015; Evers et al., 2016; Smith et al., 2017). One of the main applications of this technology has consisted in probing each gene’s potential in selectively reducing the viability of cancer cells upon inactivation (Hart et al., 2015; Tzelepis et al., 2016; Wang et al., 2014, 2015), and large panels of immortalised tumour cell lines have been CRISPR-screened with the aim of identifying genomic-context-specific cancer vulnerabilities that might be exploited therapeutically (Behan et al., 2019; Dempster et al., 2019b; Kurata et al., 2018; Martinez-Lage et al., 2018; Tsherniak et al., 2017). Other important uses of such screens have allowed to functionally characterise genes of interest (Shalem et al., 2015; Zhou et al., 2014), to identify genes that are required for cellular survival invariantly across tissues and conditions (Hart et al., 2015; Sharma et al., 2020; Vinceti et al., 2021), and to unveil novel therapeutic targets (Yu et al., 2022; Zeng et al., 2022). Beyond identifying gene essentiality and functions, CRISPR-Cas9 screens have also been used to dissect non-coding sequences and characterise regulatory and enhancer elements (Korkmaz et al., 2016; Rajagopal et al., 2016).

In a genome-wide CRISPR-Cas9 experiment, the screened models are engineered to achieve a constitutive or transient expression of the Cas9 endonuclease, and they are then transfected with a library of pooled single-guide RNAs (sgRNAs) targeting individual genes, at a genome scale.

A typical library contains multiple sgRNAs - usually from 2 to 10 (Doench et al., 2016; Gonçalves et al., 2021; Hart et al., 2015, 2017; Sanjana et al., 2014; Sanson et al., 2018; Shalem et al., 2014; Tzelepis et al., 2016; Wang et al., 2014) - targeting the same gene in different regions, in order to increase inactivation efficiency and reduce possible topological biases (Haeussler et al., 2016). In addition, each sgRNA is co-delivered together with a resistance gene to a drug-selectable marker, allowing selection of successfully transfected cells, and a tag sequence allowing counting the number of cells to which an individual sgRNA has been successfully delivered via next generation sequencing.

After selection, expansion, and genome sequencing of the transfected pool of cells, gene essentiality is usually quantified by differential representation analysis of the targeting sgRNAs. The counts derived from the DNA harvested at the end of the assay are contrasted against a control - for example, the counts derived from the transfected plasmidic DNA (Doench, 2018; Imkeller et al., 2020) - and quantified through a depletion log fold-change (logFC).

The efficiency of the CRISPR-Cas9 system originates from its mode-of-action: the induction of DNA double strand breaks (DSBs) inflicted by the Cas9 enzyme on the genomic region matched by a given sgRNA (Hsu et al., 2014). DSBs are repaired by non-homologous end joining: an error prone mechanism causing small insertions and deletions, resulting in premature stop codons, thus efficient gene silencing (Davis and Chen, 2013; Gomez and Hergovich, 2016; Panier and Durocher, 2013; Rouet et al., 1994; Shen et al., 2018; Symington and Gautier, 2011). One of the major downsides of this system is that, when used to target genomic copy number (CN) amplified regions, the Cas9 enzyme may cause a large number of DSBs. This results in a highly cytotoxic effect that is independent from the targeted gene’s function or its expression, and leads to false positive essential gene calls (Aguirre et al., 2016; Dempster et al., 2021; Gonçalves et al., 2019; Iorio et al., 2018; Meyers et al., 2017; Munoz et al., 2016; Wang et al., 2015).

A number of computational methods have been proposed by us and others to address this problem *in-silico* from the analysis of sgRNA counts and logFCs (Gonçalves et al., 2019; Iorio et al., 2018; Meyers et al., 2017; de Weck et al., 2018). We developed *CRISPRcleanR* (Iorio et al., 2018): the first tool working in an unsupervised way, that is without requiring in input any information on the copy number-alteration profiles of the screened models, and not making any initial assumption on the topological properties of the genome to which the gene-independent responses to CRISPR-Cas9 targeting might be due. CRISPRcleanR is implemented as an R (https://github.com/francescojm/CRISPRcleanR) and Python package (https://github.com/cancerit/pyCRISPRcleanR) and it is available as an image for docker and cloud environments (https://dockstore.org/containers/quay.io/wtsicgp/dockstore-pycrisprcleanr). We have used this tool in the past, upstream of bioinformatics pipelines for the identification of cancer vulnerabilities, and to perform rigorous quality control assessments of screens and their technical replicates (Behan et al., 2019; Dempster et al., 2019b). Once depletion logFCs are corrected by CRISPRcleanR, they can be scaled (Meyers et al., 2017) or normalised (Gonçalves et al., 2020) for the sake of interpretability and inter-screen comparability and/or further processed with a set of tools like BAGEL (Hart and Moffat, 2016; Kim and Hart, 2021) and MAGeCK (Li et al., 2014, 2015) to identify significantly essential genes.

The core algorithm of CRISPRcleanR operates in two steps: it first identifies genomic regions with biased sgRNA logFCs - using a circular segmentation method (Olshen et al., 2004; Venkatraman and Olshen, 2007) - then it corrects the depletion logFCs of all the sgRNAs whose targeted sequence falls in such regions. The effectiveness of CRISPRcleanR, combined with its simplicity and easily interpretable output, have made it a widely used tool (Allen et al., 2019; Chai et al., 2020; Dede et al., 2020; Gonçalves et al., 2020; Kim and Hart, 2021; Lord et al., 2020; Picco et al., 2019). However, advanced use of CRISPRcleanR in its current implementations requires solid computer programming skills: this represents a limitation for the overall accessibility and effectiveness of this tool.

In order to widen the CRISPRcleanR user base, we have developed *CRISPRcleanR*^*WebApp*^: a web-based, user-friendly, and interactive application that enables accessing all CRISPRcleanR functionalities through an intuitive graphical user interface. This application provides a wrapper around the native R package, avoiding all the low-level programming language interactions, while providing the same capabilities in terms of processing and data analysis as the original package, plus novel interactive data/result exploration modalities. Finally, CRISPRcleanR^*WebApp*^ and a recent release of CRISPRcleanR (v3.0.0) can both process low level sequencing files in FASTQ / BAM format, natively supporting a larger set of CRISPR sgRNA libraries than the original package, and encompassing six of the most popular and widespread ones (Aguirre et al., 2016; Doench et al., 2016; Gonçalves et al., 2021; Meyers et al., 2017; Ong et al., 2017; Sanjana et al., 2014; Tzelepis et al., 2016; Wang et al., 2015).

Here we provide an overview of CRISPRcleanR^*WebApp*^ functionalities and data exploration modalities. Furthermore, we report results from a comparison of CRISPRcleanR^*WebApp*^ outcomes obtained by applying it to data derived from CRISPR-Cas9 screens of the same cell line performed using our supported libraries. A high level of concordance across these outcomes indicates excellent compatibility of CRISPRcleanR^*WebApp*^ across all supported libraries.

## Design

### Overview

CRISPRcleanR^WebApp^ is a client-server web app (**Fig. 1A**), designed to use an underlying recent release of the CRISPRcleanR R package - v3.0.0, with v0.5.0 originally presented in (Iorio et al., 2018) - through a user-friendly interactive browser interface. CRISPRcleanR^WebApp^ provides a complete user experience offering the same analytical possibilities of the native version of CRISPRcleanR in terms of workflows’ usage and interactions, while enriching it with novel and interactive data exploration modalities, and the possibility of processing low level sequencing files (in FASTQ / BAM format). It consists of a web browser client Single Page Application (SPA) for user interaction, plus a backend providing data storage and processing. It also includes a user authentication/authorization mechanism, implemented through a login system, to protect submitted data and related results, ensuring a high level of privacy (STAR *Methods*). A set of video tutorials are included in the homepage of CRISPRcleanR^WebApp^, guiding the user through every step: from job submission, to results identification and access, up to results exploration and download. In addition, screen responsiveness is ensured, allowing a pleasant and effective user interaction on a wide variety of screen sizes, including tablets and smartphones. The application is served through a containerized architecture hosted at Human Technopole (the hosting research institute). This makes CRISPRcleanR^WebApp^ highly stable, easy to maintain and scalable.

**Fig. 1.**
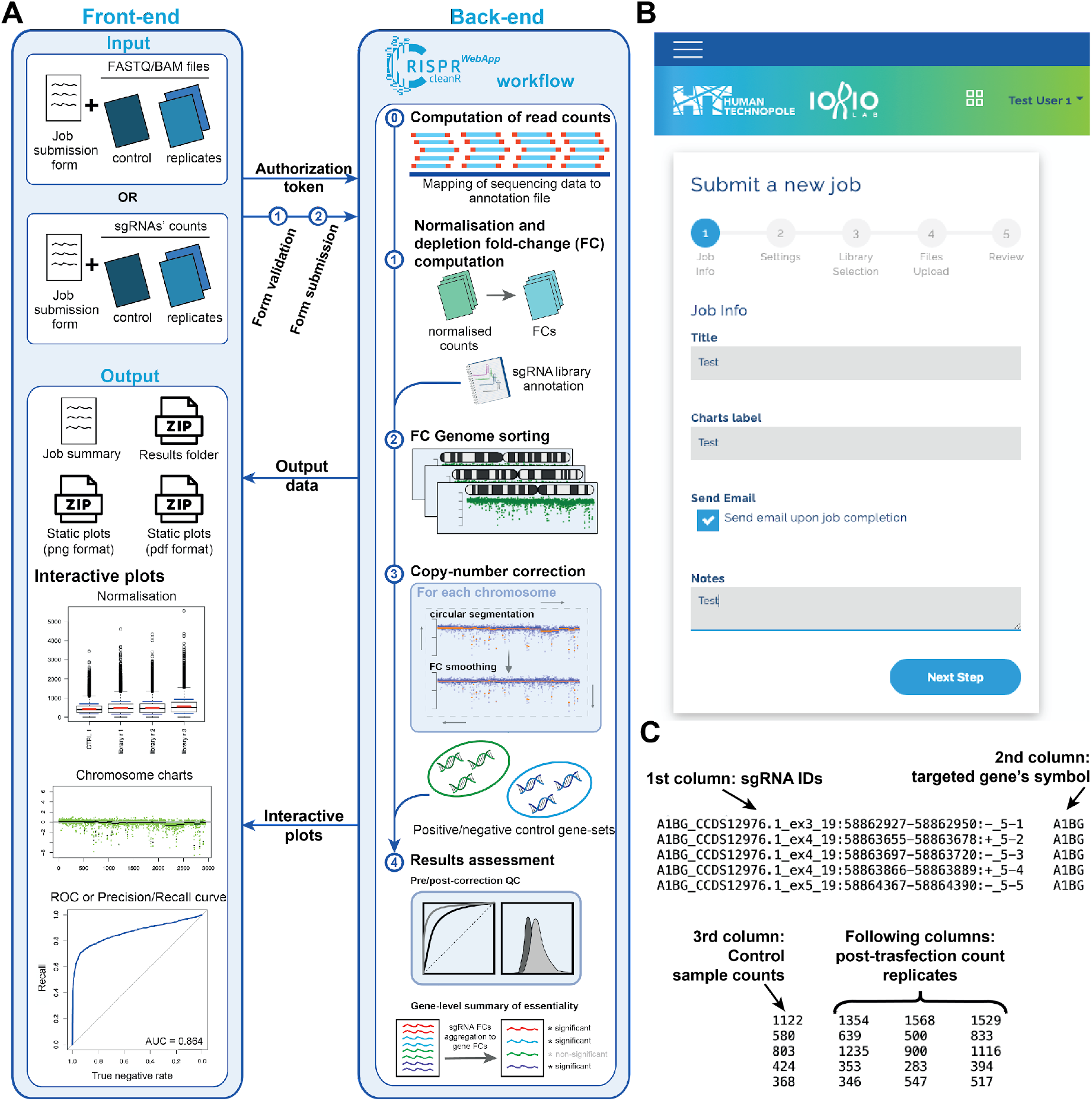
Overview of CRISPRcleanR^WebApp^ design. **A**. Schematic of the CRISPRcleanR^WebApp^ architecture. In the front-end, the user fills out a job submission form and uploads input files in FASTQ/BAM formats or pre-computed single-guide RNA read counts derived from a genome-wide CRISPR-Cas9 screen, alongside experiment metadata and library specification. The form is then validated and submitted to the backend, where the data is processed following the CRISPRcleanR workflow. Results are then made available to the web interface and they are explorable through a set of interactive plots in a dedicated results page. **B**. Step 1 of the CRISPRcleanR^WebApp^ job submission form: the entry point for starting any new job request, after secure login. As illustrated, there are fields which the user is asked to fill out before submitting the job. **C**. Example of tab-separated file containing single-guide RNA pre-computed counts. This file is derived from screening the HT-29 cell line with the Sanger KY library. First column contains sgRNA unique identifiers, the second one targeted gene symbols, then counts of the plasmidic DNA, followed by counts obtained after 14 days post-transfection and selection of the library, in three replicates.

The core function of CRISPRcleanR^WebApp^ applies a circular binary segmentation algorithm (Olshen et al., 2004; Venkatraman and Olshen, 2007) to patterns of sgRNAs’ depletion log fold-changes (logFCs) on a per chromosome basis. More specifically, it identifies genomic regions containing sgRNA clusters with sufficiently similar depletion logFCs which are, on average, significantly different from those in the flanking genomic regions. Since it is very unlikely to observe the same fitness effect when targeting a large number of contiguous genes, the logFCs in regions characterised by a stretch of essential genes (at least 3 in the default settings) are deemed as biased due to local features of the genomic segment (e.g. copy number amplification), and they are thus corrected via mean-centering. On top of that, CRISPRcleanR^WebApp^ includes a suite of tools to 1) measure, assess and visualise the effect of said correction, 2) assemble gene level summaries of essentiality and related significance scores (collapsing sgRNA logFCs by averaging on a targeted gene basis), and 3) to assess the performances of a depletion logFC rank-based classification of prior known sets of essential/nonessential genes pre/post correction. This latter classifier is also used by CRISPRcleanR^WebApp^ to identify and output genes significantly depleted at a fixed 5% false discovery rate of prior known non-essential genes, using the approach we introduced and used in (Dempster et al., 2019a; Pacini et al., 2021). Finally, CRISPRcleanR^WebApp^ implements an inverse transformation function through which corrected sgRNA counts can be derived from corrected depletion logFCs. These corrected sgRNAs are then used by CRISPRcleanR^WebApp^ to compute further summaries of gene essentiality, and associated significance scores, via mean-variance modelling (using MAGeCK (Li et al., 2014)). Taken together, these features make CRISPRcleanR^WebApp^ a one-stop tool for the complete processing and analysis of CRISPR-Cas9 screens, producing results that are readily usable and interpretable by non computational scientists. These results can also be post-processed by bioinformaticians to call depletion significance in a supervised manner, employing classification templates (Vinceti et al., 2022) via Bayesian statistics, using BAGEL (Hart and Moffat, 2016; Kim and Hart, 2021).

### Interface and workflow setup

CRISPRcleanR^WebApp^ implements two main analytical workflows: the first one is for preprocessing input files (i.e. raw sgRNA counts as FASTQ, BAM, or pre-computed in text format) as well as normalising and correcting sgRNAs counts and depletion logFCs; the second one implements a series of data quality control (QC) assessments, and outputs interactive visualisations and result files. The first workflow encompasses calls to the complete set of CRISPRcleanR functions.

FASTQ files, are converted to sgRNA counts by using the mapping sequences included in the sgRNA library annotation (derived either from one of the CRISPRcleanR built-in data object or from a plain text file provided by the user, STAR *Methods*). The BAM format is also supported: in this case the sequence identifiers are mapped to the guide identifiers to generate the sgRNA counts (STAR *Methods*). The user can optionally decide to upload pre-computed sgRNA counts as plain text file.

Following this, different setups can be selected for the normalisation step: for instance, sgRNA raw counts can be normalised either by scaling sample-wise, based on the total number of reads, or via the median ratios’ method (Maza et al., 2013). In addition, the user can specify the number of control samples should this be greater than one (default value). Another option is to explicitly specify the minimal value of read counts for a sgRNA in the control sample in order to be included in the follow-up analyses (default value is 30 as in (Behan et al., 2019)). Finally the CRISPRcleanR correction is executed, using default values for the parameters of the relevant functions.

The second workflow encompasses all steps needed to assess the quality of a CRISPR-Cas9 screen and visualise the results. For instance, common quality checks are based on the computation of the Area Under Receiver Operating Characteristic and Precision-Recall curves (AUROC and AUPRC, respectively). In particular, the computed profile of gene/sgRNA depletion logFCs are employed as a rank-based classifier of two built-in sets of prior known essential and nonessential genes (or their targeting sgRNAs). All the results are then summarised in data plots that can be queried through interactive visualisations or downloaded in static graphic formats (i.e. PNG or PDF).

These workflows are seamlessly integrated within CRISPRcleanR^WebApp^. Indeed, after secure login, the user only needs to fill out the initial parameters, including a minimal amount of experimental metadata, in a job submission form (**Fig. 1B** and **Table 1**), and upload properly formatted input files containing sequencing data (FASTQ/BAM files) or the sgRNA pre-computed counts (**Fig. 1C**) to run the analyses.

**Table 1.** Fields to be filled in by the user in the job submission form.

| Submission form step | Field | Type | Mandatory ? | Description |
| --- | --- | --- | --- | --- |
| **1**<br>**(Job Info)** | Title | String | Yes | Used to identify the output in the results section once the job is completed. |
|  | Charts label | String | Yes | A string that will be used as a title for all the plots generated by CRISPRcleanR. |
|  | Send Email | Checkbox | Optional | If selected the user will receive a notification email upon job completion. |
|  | Notes | String | Optional | Additional notes describing the job |
| **2**<br>**(Settings)** | Minimal n° reads in the control sample | Integer | Yes (default value = 30) | Minimal n° reads that each sgRNA should have in the control sample (or on average across control samples) in order to be included in the analysis. |
|  | Normalisation method | Multiple selection | Yes (default value = Scaling by total numbers of reads) | Normalisation method for the read counts (scaling by total numbers of reads or Median Ratios). |
| **3**<br>**(Library Selection)** | Library Type | Mutually exclusive radio buttons | Yes (default value = Built-in) | Switch between one of the natively supported sgRNA libraries or other (external) library. |
|  | Built-in library | Multiple selection | Yes if the selected library type is 'Built in'; inactive otherwise | Allows selecting one of the natively supported sgRNA libraries (AVANA, Brunello, GeCKO, KY v1.0, KY v1.1, MiniLibCas9 and Whitehead). |
|  | Library annotation file | File upload | Yes if the selected library type is 'Other'; inactive otherwise | Allows uploading a library annotation file. |
| **4**<br>**(Files upload)** | Data type | Mutually exclusive radio buttons | Yes (default value = sgRNA counts) | Allows switching between pre-computed sgRNA counts (in plain text format), FASTQ or BAM files. |
|  | sgRNA counts file | File upload | Yes if the selected Data type is 'sgRNA counts'; inactive otherwise | Allows uploading pre-computed sgRNA counts as plain text file. |
|  | N° of controls | Integer | Yes with default value = 1 (if selected Data type is 'sgRNA counts'; inactive otherwise) | N° of control samples in the screen, i.e. columns in the sgRNA count file. |
|  | FASTQ controls | File upload | Yes if the selected Data type is 'FASTQ'; inactive otherwise | Allows uploading control sample(s) as FASTQ files. |
|  | FASTQ samples | File upload | Yes if the selected Data type is 'FASTQ'; inactive otherwise | Allows uploading screen replicates as FASTQ files. |
|  | BAM controls | File upload | Yes if the selected Data type is 'BAM'; inactive otherwise | Allows uploading control sample(s) as BAM files. |
|  | BAM samples | File upload | Yes if the selected Data type is 'BAM'; inactive otherwise | Allows uploading screen replicates as BAM files. |
| **5**<br>**(Review)** | Submit | Button | Yes | Job submission finalisation. |

If the user opts to upload pre-computed read counts, these need to be in a comma- or tab-separated plain text with csv and tsv extensions. In this file, the first two columns must include unique sgRNA identifiers and their targeted gene’s symbol, respectively, followed by one or more (in case of multiple controls) columns containing plasmid/control read counts. The remaining columns must contain replicates of the sgRNAs’ counts obtained post-selection and amplification, in the CRISPR screen (**Fig. 1C**). Before submission, the entire form is checked for potentially incorrect file formats or missing parameter specifications, in order to prevent incomplete or inconsistent input data.

After job submission, the user is notified of the outcome of this process (i.e. successful submission or submission error).

The CRISPRcleanR WebApp home page contains a link to download example input files in a single compressed folder. This folder contains pre-computed sgRNAs’ read counts obtained by screening the HT-29 cell line with the Sanger KY library (Tzelepis et al., 2016) in a study by Behan et al (Behan et al., 2019), and it includes one control sample, i.e. read counts from the plasmid DNA, and three post-selection/amplification read counts’ replicates (fourth, fifth and sixth columns). Read counts from a similar experiment are also included in FASTQ format (one for the plasmid/control DNA, i.e. test_plasmid.fq.gz, and two for the screen replicates, i.e. test_sample1.fq.gz and test_sample2.fq.gz, downsampled to reduce file size). Finally, the example data folder contains a text file with the annotation of the KY v1.0 library and a readMe file describing the content of the folder.

### Results exploration and interactive plots

Once a job has been submitted, the CRISPRcleanR^WebApp^ back-end server starts an offline computation, sending an email message to the user once finished (if this option was selected in the job submission form). A new results’ entry is then immediately listed in the results page (**Fig. 2A**), where a data table offers a configurable pagination size for splitting large sets of entries into smaller chunks. In this table, each row refers to a job submitted by the logged user, and it shows main details such as submission date and time, job title, and job status (succeed, failed or pending). Furthermore, the table allows for jobs filtering and sorting according to any of the column fields. Through this page the user can access and interactively explore the results obtained from each of their job submissions (**Fig. 2B**). The job results page shows a detailed summary that recapitulates the parameters specified by the user in the job submission form, and a series of panels allowing to explore and download all data and results, and to access all the interactive plots outputted by CRISPRcleanR^WebApp^. The Downloads panel contains links to all the results, which can be downloaded as zipped folders, as well as to all the plots, downloadable as static images in pdf or png format.

**Fig. 2.**
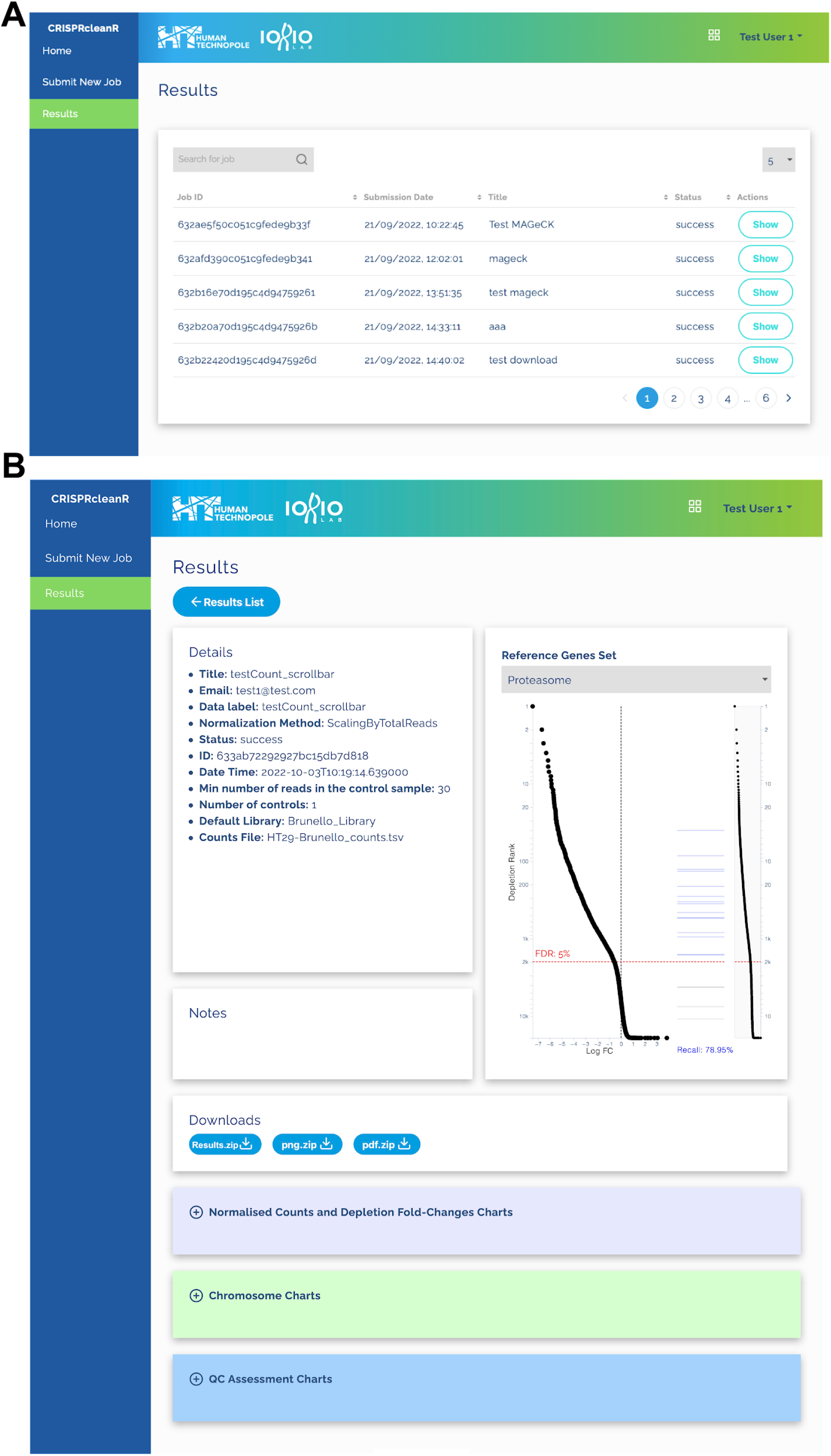
Results section of CRISPRcleanR^WebApp^. **A**. The results list page: here the user will find all the results obtained from their submitted jobs. **B**. Job results accessible by selecting a job ID in the results list page. The page is made of several sections including a detailed description of the job, an interactive summary plot where all the screened genes are ranked based on their depletion fold-changes with overlaid signatures or control genes. Three image accordions contain clickable thumbnails for opening the related interactive plot rendered by the web application. Finally, the Downloads section includes links to all plots as static images (in pdf or png format), as well as to both input and processed data and results (as unique zipped folder of plain text files).

All the plots can be also visualised interactively and are equipped with a tooltip providing detailed information when hovering the pointer on a graphic component. The job results page includes also a summary *gene-signature* plot, while the other plots are collected within image accordions, containing clickable thumbnails, and partitioned in three different panels (**Fig. 2B**): normalised counts and depletion fold-changes charts, chromosome charts and QC assessment charts. All charts include a zoom area, often showing a minigraph representation of the overall chart, where users can select an area to be magnified in the main chart. The gene-signature plot interactively shows results from the normalisation and the depletion logFCs and count correction, i.e. the *chromosome plots*. On the top-left of the results’ page is provided a plot of all screened genes with coordinates indicating depletion logFCs (x-axis) and depletion rank position (y-axis), respectively (**Fig. 2B**); on the top, the user can select one among 7 different signatures of control genes, i.e. prior known essential gene sets like proteasome, spliceosome, DNA replication, ribosomal proteins, RNA polymerase, and BAGEL essential and nonessential gene-sets (Hart et al., 2014, 2017)), and explore how they rank based on their depletion logFCs, on the right part of the chart. Apart from the BAGEL gene-sets, the other 5 signatures are derived from the Molecular Signature Database (MSigDB) (Iorio et al., 2018; Subramanian et al., 2005). The red line indicates the rank position above which a false discovery rate (FDR) of non-essential genes is < 5% (when considering all the genes in previous rank positions as positive predictions) and it is determined using the logFC distributions of the BAGEL essential and nonessential genes (STAR *Methods*). The “normalised counts and depletion fold-changes” panel contains two interactive plots: the first one shows a comparison between raw and normalised sgRNA read counts (**Fig. 3A**), whereas the second shows uncorrected depletion logFCs (**Fig. 3C**), across samples in the input file.

**Fig. 3.**
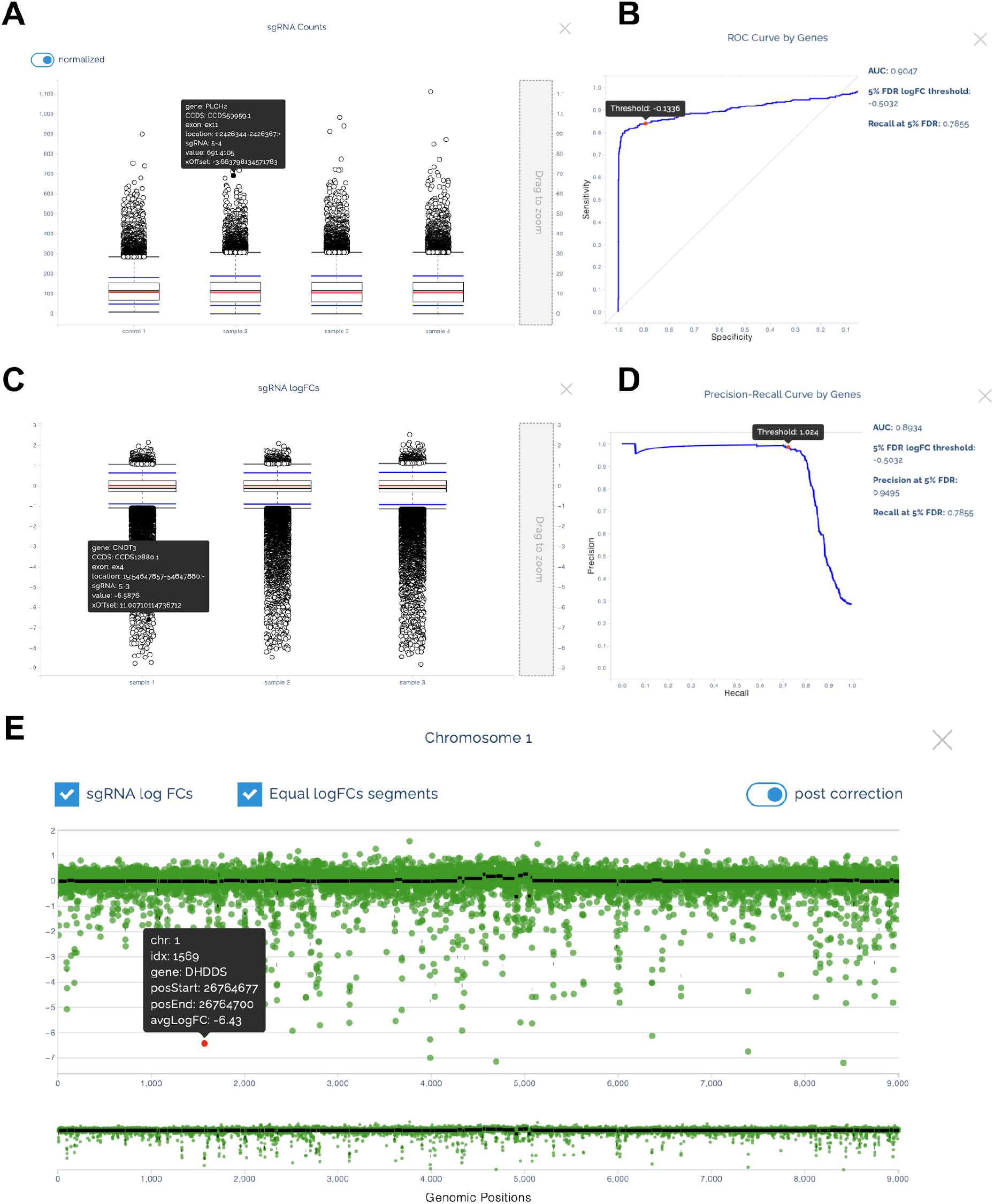
Interactive plots rendered by CRISPRcleanR^WebApp^. **AC**. Overview of the normalisation plots: **A**. boxplots for raw and normalised single-guide RNA (sgRNA) read counts across samples in the input file. **C**. CRISPRcleanR uncorrected fold-changes (FCs). Both plots have a vertical scrollbar on the right that allows zooming in on specific portions of the plot. When overing on each point, a tooltip shows information regarding the corresponding sgRNA, i.e. guide ID, exon, gene, and read count/ depletion logFC. **BD**. Quality control (QC) assessment plots. **BD**. Examples of interactive Receiver Operating Characteristic (ROC) and Precision Recall Curve (PRC) plots, obtained from profiles of corrected depletion logFCs when considering them as rank-based classifiers of two sets of a negative/positive control genes, i.e. a priori known essential and nonessential genes. **E**. Interactive plot showing the sgRNA logFCs (green dots) obtained targeting genes in the chromosome 1. The logFCs are clustered within segments of equal copy number (black lines), which are corrected by CRISPRcleanR. Each point is a sgRNA and when moving the mouse over it a tooltip will show related information such as chromosome number, guide ID, start and end position and depletion logFC when hovering the pointer on it.

The user can toggle between raw and normalised read counts by clicking on a switch button in the upper left corner of the corresponding interactive plot (**Fig. 3A**). A tooltip provides several information on a given point, i.e. guide ID, exon, gene, and raw or normalised read count, when moving the mouse over it. The same functionalities, with the exception of the switch button, are accessible in the interactive plot showing the uncorrected logFCs (**Fig. 3C**).

The “QC assessment” panel contains a series of plots summarising data quality checks performed on the sgRNA/gene-level depletion logFCs corrected by CRISPRcleanR^WebApp^ (**Fig. 3BD**). Particularly, they include visualisations of ROC and Precision/Recall curves (**Fig. 3BD**) computed on sgRNA- or gene-level corrected depletion logFC profiles, when they are considered as rank-based classifiers of two built-in reference sets of a priori known essential and nonessential genes. A tooltip provides information regarding the logFC threshold below which a certain level of recall (for the AUROC plot), or precision (for the AUPRC plot) is achieved.

The “chromosome charts’’ panel shows one plot per chromosome, with segments of sufficiently similar sgRNAs’ depletion logFCs on the x-axis, and the corresponding average logFC pre-/post-CRISPCcleanR-correction on the y-axis. The first 22 charts correspond to the autosomal chromosomes, the 23^rd^ chart corresponds to the X chromosome, and the 24th chart to the Y or X chromosome. Each chromosome chart has two checkboxes, in the upper left corner, allowing the user to focus just on the segments (black lines) or the sgRNA depletion logFC (green dots) components. A switch button, in the upper right corner, allows to toggle between uncorrected and corrected logFCs. Also in this case, the segment tooltip shows relevant information such as chromosome number, segment ID, start and end position of the segment location on the genome, and average depletion logFC of the mapped sgRNAs; on the other hand, the sgRNA tooltip provides information about chromosome number, targeted gene, start and end position of the gene portion targeted by the sgRNA, and depletion logFCs (uncorrected or corrected).

## Results

### Extended sgRNA libraries’ support

The original version of CRISPRcleanR (v0.5.0) supported only the AVANA (Doench et al., 2016) and Sanger KY (Behan et al., 2019; Tzelepis et al., 2016) CRISPR-Cas9 sgRNA libraries. CRISPRcleanR^WebApp^ builds upon and uses CRISPRcleanR v3.0.0, which (from v2.0.0) we have extended to fully support the following additional genome-wide libraries: Brunello (Sanson et al., 2018), GeCKOv2 (Sanjana et al., 2014; Shalem et al., 2014), Whitehead (Park et al., 2017; Wang et al., 2014, 2015), and the recent MiniLibCas9 (Gonçalves et al., 2021) library of minimal size (with only two targeting sgRNA per gene). With the exception of GeCKOv2, the (publicly available) annotations of all these libraries include genomic coordinates of the targeted sequences for all the sgRNAs. This information is needed by CRISPRcleanR to genome-sort sgRNA depletion logFCs prior correction. For the GeCKOv2 library, this information was not available. For this reason we remapped the GeCKOv2 sgRNA sequences onto the human genome (GRCh38 - hg38) and assembled a CRISPRcleanR compatible annotation object (STAR *Methods*) for this library. In addition, CRISPRcleanR^WebApp^ supports any genome-wide CRISPR library provided that the user uploads a custom library annotation file. In this case, the sgRNA sequences in the read count files are matched to the ones provided in the library annotation file, and the workflow proceeds in case of a successful retrieval of at least 80% of the guides (STAR *Methods*). This functionality further extends the applicability of CRISPRcleanR^WebApp^ to a larger pool of CRISPR screens performed on 2D cell lines or alternative cancer models (e.g. primary cultures, organoids or patient-derived xenografts).

We tested the ability of CRISPRcleanR in supporting the extended set of libraries described above, in terms of correction performances and results’ conservation across analyses of screens performed on the same cell line. Particularly, we tested CRISPRcleanR on screens performed with all the aforementioned genome-wide CRISPR-Cas9 libraries on the HT-29 cell line. HT-29 is a human cancer cell line derived from colorectal carcinoma frequently used to assess sensitivity and specificity of CRISPR-Cas9 libraries and with multiple publicly available CRISPR screen datasets (Behan et al., 2019; Doench et al., 2016).

First, we compared the extent of correction brought about by CRISPRcleanR on the different screens. To this aim we contrasted gene depletion logFCs pre-versus post-correction, and compared differences across libraries (**Fig. 4A**). Prior to this comparison we averaged sgRNA-level depletion logFCs on a targeted gene basis, in order to obtain gene-level depletion logFCs. As the tested screens presented different depletion logFC ranges and phenotype penetrance, due to inherent differences in the employed sgRNAs’ sequences and for the sake of inter-screen comparability, we concatenated screen-wise the pre- and post-correction logFC vectors and applied a min-max normalisation (STAR *Methods*) before comparing the different screens. After the normalisation, we split back each normalised vector in the two original components and computed differences between pre- and post-corrected logFCs.

**Fig. 4.**
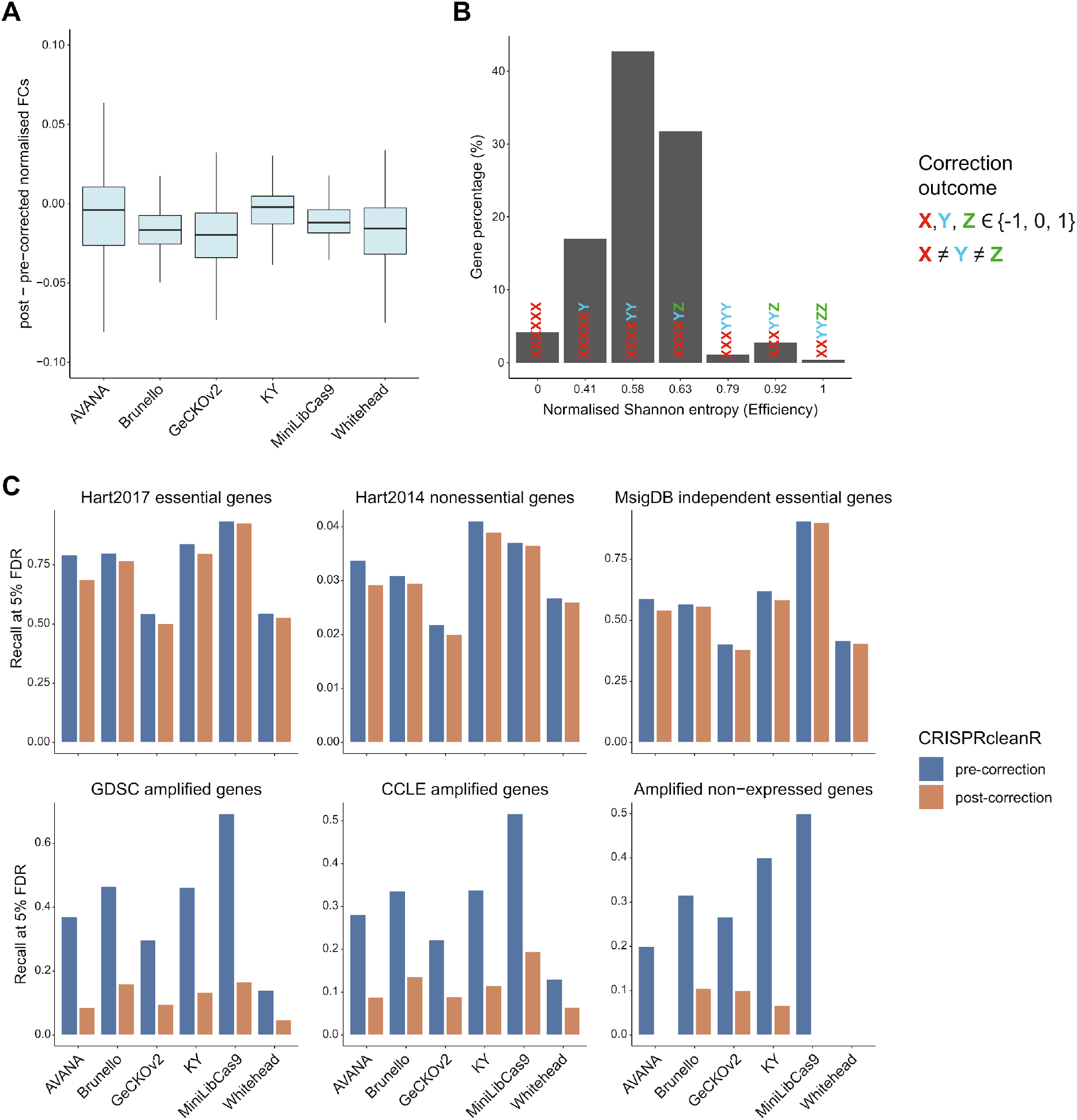
Assessment of CRISPRcleanR correction across different supported CRISPR-Cas9 libraries on the HT-29 cell line. **A**. Comparison of CRISPRcleanR pre- and post-correction fold-changes (FCs) across screens. **B**. Normalised Shannon entropy (efficiency) of gene-wise correction outcomes (−1, 0, 1) across screens. Only genes shared across libraries were considered. **C**. Recall at 5% FDR of six prior known gene-sets observed when considering pre- and post-correction (as indicated by the different colours) sgRNA logFCs’ profiles as rank-based classifiers.

We observed an average median of these logFC differences across library equal to −0.012 (min = −0.002 for KY and max = −0.02 for GeCKOv2) and an average interquartile range equal to 0.0245 (min = 0.015 for MiniLibCas9 and max = 0.037 for AVANA (**Fig. 4A**). These results show that the correction applied by CRISPRcleanR has limited effects on the whole screen, it is focused on a small set of genes and that this minimal impact is conserved across screens of the same cell line performed with the different supported libraries.

Next, we asked at what extent the CRISPRcleanR correction affected each gene’s logFC consistently across screens. To this aim, we computed for each gene a normalised Shannon entropy, also known as efficiency, quantifying how homogeneous were the correction effects for a given gene across screens. A low entropy value indicated that a gene’s logFC was affected consistently across screens, whereas a high entropy value indicated the opposite, (STAR *Methods*). In particular, we coded the gene-wise correction outcomes in a given screen as follows. A 0 indicated that the logFC of the gene under consideration was not corrected by CRISPRcleanR, as the sgRNAs targeting that gene were not mapped onto a genomic segment detected as biased. A 1 indicated a positive correction, meaning that the sgRNAs targeting the gene under consideration were mapped onto a genomic segment detected as biased toward negative values by CRISPRcleanR and their logFC increased. Following the same logic, a −1 indicated a negative correction. Applying this coding across all screens, yielded for each gene a vector of 6 entries (one per each tested library) with values in {-1, 0, 1}, from which we computed the normalised Shannon entropy.

A summary of the results is provided in **Fig. 4B**, showing the percentages of genes across efficiency values and concordance/discordance of correction effects for each value. For example, the first bar from the left (corresponding to a 0 efficiency) accounted for all the genes for which the correction outcomes were identical across screens, whereas the second bar accounts for the corrections that agree across all the screen but one, and so on. For over 95% (**Fig. 4B**) of the genes we observed the same correction effect in at least four screens out of six (efficiency < 0.63), with the absolute majority (42%) showing different outcomes in a 4:2 proportion. Finally, only 4.31% of the genes presented all three different correction outcomes across screens, and 0.39% presented them equally partitioned (i.e. two −1s, two 0s, and two 1s). Overall these results show that the CRISPRcleanR correction affected individual genes’ logFCs homogeneously across screens of the same cell line performed with the different supported libraries.

Finally, we measured the extent to which the CRISPRcleanR correction tendency in reducing false positive gene-essentiality calls while maintaining true positive calls was conserved across processed screens.

To this aim, we considered six control sets of genes (**Table 2**): the Hart2017 essentials (Hart et al., 2017), Hart2014 nonessentials (Hart et al., 2014), respectively as positive and negative control plus another set of known essential gene-sets derived from the MSigDB (Pacini et al., 2021; Subramanian et al., 2005) accounting for housekeeping cellular processes (DNA replication, histone genes, RNA polymerase, proteasome, ribosomal protein genes and spliceosome), and finally two sets of HT29 specific putative negative controls, i.e. copy number amplified genes (from two different public resources) (Barretina et al., 2012; Iorio et al., 2016; Mermel et al., 2011), as well as amplified and non-expressed genes (FPKM < 0.1 in HT-29, STAR *Methods*).

**Table 2.** Sets of predefined genes for the assessment of CRISPRcleanR correction performances.

| Set name | Set Type | Number of genes | Dataset of origin |
| --- | --- | --- | --- |
| Hart2017 essential | Reference set of core-fitness essential genes. | 684 | BAGEL reanalysis of 17 genome-scale knockout screens in human cell lines performed with different libraries (Hart et al., 2017) |
| Hart2014 nonessential | Reference set of nonessential genes. | 927 | Large collection of shRNA gene dependency profiles analysed with a linear algebra approach (Hart et al., 2014). |
| MSigDB independent essential | Pooled sets of independent essential genes from the MSigDB. | 290 | MSigDB dataset (Pacini et al., 2021; Subramanian et al., 2005) |
| GDSC amplified | Amplified genes in HT-29 derived from the GDSC dataset. | 105 | GDSC dataset (Iorio et al., 2016). |
| CCLE amplified | Amplified genes in HT-29 derived from CCLE dataset. | 198 | CCLE dataset (Barretina et al., 2012; Mermel et al., 2011). |
| Amplified non-expressed | Amplified non-expressed (FPKM < 0.1) genes in HT-29 derived from the GDSC dataset. | 21 | GDSC dataset (Iorio et al., 2016). |

For each gene-set, we computed the recall at 5% FDR (Star *Methods*) for the pre- and post-corrected logFCs across the six screens (**Fig. 4C**). For the negative controls (expected to be biased) we observed consistent reductions in recall across screens (median = 29.48%, 19.65% and 20.53%, respectively). In contrast, for the positive controls, we observed negligible recall reductions across screens (median = of 3.61%, 0.16% and 1.74% respectively). Thus, CRISPRcleanR can effectively correct the logFCs of amplified gene-sets with high specificity without compromising the logFCs of other genes, regardless of their phenotype intensity and employed supported library.

## Discussion

We introduced CRISPRcleanR^WebApp^, a web-based application integrating the complete suite of functionalities available in the CRISPRcleanR R/python package (Iorio et al., 2018). CRISPRcleanR is a computational tool for correcting gene-independent responses to CRISPR-Cas9 targeting that are observed in data from pooled viability screens and arise from copy number amplifications. Differently from other methods, CRISPRcleanR works in an unsupervised way not requiring any input related to the copy number variations profiles of the processed/screened model. In addition, CRISPRcleanR carries out the correction on a single-sample basis, not requiring multiple screens to be analysed jointly nor borrowing signals across screens. This offers the benefit of preserving the overall heterogeneity of the data, making our tool especially suited for the identification of context-specific dependencies and biomarkers (Pacini et al., 2021).

We also showed that CRISPRcleanR^WebApp^ does not require any prior knowledge of programming languages like R and python, and offers a user-friendly interface giving full workflow control, and fully customisable execution of the correction procedure on the data provided by the user.

The homepage of CRISPRcleanR^WebApp^ is equipped with comprehensive video-tutorials on its usage and a toy dataset for testing. The job submission form can take in input FASTQ/BAM files (which are subject to quality checks and are then mapped to the library annotation file to obtain sgRNA counts), or pre-computed sgRNA counts in a plain text format. A ‘Results’ page allows users to download the output data, such as corrected logFC file/s and static plots, as well as exploring the results through a portfolio of interactive plots.

Furthermore, CRISPRcleanR^WebApp^ supports a larger set of built-in genome-wide CRISPR libraries compared to the original R package. We also enabled users to upload data from screens performed with custom libraries: in this case, sgRNA IDs are checked for consistency with respect to the library annotation file. We believe this feature will extend the services of CRISPRcleanR^WebApp^ to a much larger audience. Indeed, CRISPR-Cas9 screens can be carried out in a host of different models besides 2D cell lines (e.g. primary cultures, organoids, or patient-derived xenografts). These screens are amenable to CRISPRcleanR correction, provided that they are performed using libraries with sufficient sgRNA density, and related annotation available.

Here, we have provided an overview of the CRISPRcleanR^WebApp^ implementation, design and interface and demonstrated that it yields consistent results across different technical settings and supported libraries. Taken together, the features of CRISPRcleanR^WebApp^ provide an easy-to-use framework for pre-processing and correcting data derived from CRISPR-Cas9 screens, which might significantly widen the CRISPRcleanR user-community.

### Limitations of the study

While CRISPRcleanR^WebApp^ is a user-friendly web application accessible to non-computational scientists, its current version still lacks some functionality of the original package, which will be included in subsequent versions. For instance, we are planning to extend the number of input parameters that can be specified before submitting a job, as experienced users may require more advanced setups.

In addition, CRISPRcleanR is equipped with functions to test the depletion logFCs of sgRNAs targeting different reference gene sets (for example prior known essential genes, or copy number amplified genes) for statistically significant differences with respect to the background pre- and post-CRISPRcleanR correction. The aim of this analysis is to show that the CRISPRcleanR correction reduces false-positive essential gene calls while maintaining true-positive rates. These functions will be visually rendered in future versions of CRISPRcleanR^WebApp^. Furthermore, CRISPRcleanR includes the ccr.impactOnPhenotype function that computes the percentages of genes whose depletion signal is attenuated post-CRISPRcleanR correction or potentially ‘distorted’ (i.e. loss-of-fitness genes in the uncorrected screens becoming gain-of-fitness genes post-correction, and vice-versa). In (Iorio et al., 2018) we demonstrated that the amount of this type of ‘distortion’ introduced by CRISPRcleanR is negligible, however also this analysis will be possible in future versions of the CRISPRcleanR^WebApp^. Finally, another feature we are planning to integrate in new versions is the possibility to pipeline CRISPRcleanR^WebApp^ with existing tools that robustly estimate gene essentiality after depletion logFC correction supervisedly (like BAGEL (Hart and Moffat, 2016; Kim and Hart, 2021).

## Methods

### Web-application architecture

#### Front-end

The front-end web application (also known as client) is implemented as a Single Page Application (SPA). Dynamic data is retrieved from a backend API server, returning data in JSON format, which is then used from the SPA to render HTML/CSS accordingly. Example files are made downloadable from the API server, while all job related files are managed through a dedicated file server.

The client is based on Vue.js JavaScript framework (https://vuejs.org/). Interactive charts are implemented as scalable vector graphics documents managed through Vue components leveraging the D3.js library (https://d3js.org/). Design and styling were done entirely from scratch with Sass stylesheets, compiled as CSS during the application bundling process. Screen layout and responsiveness is achieved with Flexbox, CSS Grid layouts and Javascript. The bundled web application is served through a Dockerized (https://www.docker.com/) version of Nginx (https://www.nginx.com/), hosted on a virtual machine within HT IT infrastructure.

#### Back-end

The back-end consists of a docker multi-container app built upon the following containers and underlying technologies:

- Application Programming Interface (API) server: FastAPI python framework (https://fastapi.tiangolo.com/)
- File server: NodeJS with Express.js framework
- Background queue: Celery Python (https://docs.celeryproject.org/en/stable/getting-started/introduction.html)
- Message Broker: Redis (https://redis.io/)
- Database: MongoDB (https://www.mongodb.com/)

An on premises S3 bucket compliant object storage is used to store all job related files (both input and output).

##### Application Programming Interface

The API server is implemented with FastAPI, a modern and popular Python API framework.

It is provided as a Docker container based on a python bullseye official docker image.

##### File server

A NodeJS server is used to manage the files’ upload and download. This allows to directly and reliably stream data files to and from the S3 bucket, since it is not possible with FastAPI. A redis-based pub-sub messaging system exists to ensure the file server informs back the API main server about the upload outcome.

##### Task queue and Message Broker

To maintain the backend API server responsive while processing a job, we implemented an offline job processing mechanism. More specifically, job processing is delegated to another container that receives jobs and processes them asynchronously with respect to the actual job submission, following a producer-consumer pattern. When the backend receives a new submitted job, this is forwarded to the background task queue through the message broker to inform consumers about new tasks to be executed.

Background processing is implemented through Celery, a common python task queue manager. Celery is executed on a dedicated Docker container based on a python bullseye official docker image, further customised to perform R processing. This customised image installs a Debian-compatible R distribution, along with all required CRISPRcleanR dependencies. Communication with CRISPRcleanR package is performed through Rpy2 (https://rpy2.github.io/), a python package that enables us to register R functions and environments into python wrapping objects, such that R interactions can be directly executed from python code.

During job processing, the background queue container instantiates a single celery consumer, called worker, that consumes job computation requests from the message queue. In order to obtain parallel processing over multiple jobs, distinct independent workers can be spawned on container replicas by properly configuring the underlying container orchestrator. The message broker is then able to automatically send new job computations according to each worker’s current workload, while still avoiding the same job execution to be performed on different workers. The communication between FastAPI backend and task queue is accomplished through a Redis-based message broker, the latter running on a dedicated container.

##### Database Container

Our database container runs on a MongoDB instance, a cross-platform document-oriented NoSQL database. In addition, the backend container communicates with MongoDB using a Motor asyncio driver (https://motor.readthedocs.io/en/stable/), whereas the Celery-based background queue uses a Pymongo driver (https://pymongo.readthedocs.io/en/stable/), not being based on asyncio patterns.

##### Security

CRISPRcleanR^WebApp^ implements an authorization schema based on OAuth2 protocol (https://datatracker.ietf.org/doc/html/rfc6749), following the Authorization Code Flow with a public client. A further layer of authentication is provided through OpenID Connect (https://openid.net/connect/), which enables us to transmit user information, such as username and email, to the client application. This highly secure schema allows for great flexibility in configuration and it might easily be extended to implement federated/third party access through external acknowledged identity providers.

The security schema is implemented through Keycloak (https://www.keycloak.org/), an integrated authorization/authentication system which provides a comprehensive solution for managing users and providing an authorization server for managing authentication and applications’ tokens. Our hosting infrastructure at Fondazione Human Technopole is equipped with a dedicated Keycloak system made accessible both from the backend (for token validation) and frontend (for issuing tokens).

### CRISPRcleanR^WebApp^ management of input files

#### Computation of read count from FASTQ or BAM files

CRISPRcleanR^WebApp^ accepts trimmed FASTQ as well as BAM file formats as input to derive single-guide RNA (sgRNA) counts. For the FASTQ format, a preliminary quality control is performed on the sequencing data based on the quality scores of the corresponding nucleotide sequences. The sgRNA sequences are then mapped to the library index, which it’s generated from the sequences retrieved in the library annotation file using the Rsubread R package (Liao et al., 2019). In order to provide the most reliable counts estimation the alignment doesn’t allow N bases, mismatches or gaps. All alignment summary statistics are provided in a text file available in the downloadable results. BAM files generated by the alignment step or supplied as input are processed using the GenomicAlignments R package (Lawrence et al., 2013). The occurrence of the “seqnames” of the aligned reads are counted and matched with the sgRNA IDs in the library annotation to provide the counts for each sample. All sample count data are then merged to create a count matrix that is downloadable from the results page and used in the following steps of the pipeline.

#### Upload of custom genome-wide CRISPR-Cas9 library

In case of upload of a custom genome-wide CRISPR library (i.e. not part of the six built-in libraries of CRISPRcleanR^WebApp^), the annotation file must include a “seq” field including the sgRNA sequences used in the screening. The application will then convert those sequences in a library index suitable for the alignment and then proceed to evaluate the read counts as described for the standard libraries. The matching has to be exact (i.e. no mismatches allowed), and at least 80% of the guides must be recapitulated to proceed with the workflow of the application and correct the depletion fold-changes for biases.

### Validation of the supported CRISPR-Cas9 libraries in the new version of CRISPRcleanR

#### Data acquisition

We validated CRISPRcleanR (v2.2.1) on six screens obtained from transducing popular CRISPR-Cas9 libraries into the HT-29 cell line. We tested the following libraries: AVANA (Doench et al., 2016), Brunello (Sanson et al., 2018), GeCKOv2 (Sanjana et al., 2014; Shalem et al., 2014), KY (Behan et al., 2019; Tzelepis et al., 2016), MiniLibCas9 (Gonçalves et al., 2021) and Whitehead (Park et al., 2017; Wang et al., 2014, 2015). All raw read count files are available in the following GitHub repository: https://github.com/francescojm/CRISPRcleanR/tree/master/inst/extdata.

#### GeCKOv2 library mapping onto the Human genome

We mapped the protospacer sequence of each sgRNA in the GeCKOv2 library onto the human genome (GRCh38 - hg38) using the short read mapping method bwa-mem. Most of the reads were mapped uniquely to the reference genome sequence and their positions were found within their targeted genes. On the other hand, the remaining guides were mapped ambiguously and with some mismatches. We remapped them to the reference human genome using first BLATt and then BLASTn. For the guides mapped onto multiple locations, we only considered those included in the targeted genes declared in the original library annotation file. The annotations of the genes were extracted from Gencode v38. Finally, the remaining sequences were mapped to the reference genome with mismatches, and the positions with the minimum mismatches within the targeted genes were selected. All the guides were mapped to a position on the reference genome within their targeted genes.

#### Comparison of CRISPRcleanR pre- and post-correction logFCs across screens

To compare the differences in log fold-changes (logFCs) pre- and post-correction, we first normalised the sgRNAs’ raw read counts by scaling for the total number of reads, after filtering out those guides having a plasmid read count less than 30. This is implemented in the *ccr*.*NormfoldChanges* function and resulted in uncorrected depletion logFCs. Following the pipeline implemented in https://github.com/francescojm/CRISPRcleanR/blob/master/Quick_start.pdf, we applied the *ccr*.*GWclean* function with default settings to correct the logFCs of each screen.

We obtained gene-level logFCs by averaging the score of sgRNAs targeting the same gene, and considering only the genes in common across the six screens, leading to 8473 genes. In order to reduce technical variability due to the usage of different libraries, we applied for each screen a min-max normalisation on the concatenated vector of pre- and post-corrected gene-level logFCs. We then split back the vector in pre- and post-corrected logFCs, and computed gene-wise logFC differences across screens.

#### Gene-wise correction agreement across screens

In information theory, Shannon entropy is used to quantify the uncertainty of a random variable. It is formally defined as the average auto information associated with each outcome of the random variable. Efficiency (or normalised Shannon Entropy) is defined as the Shannon entropy divided by the maximum value that the entropy can assume (which depends on the size of the spectrum of the random variable under consideration). In our case: the lower the efficiency the higher the homogeneity in the CRISPRcleanR outcomes across screens. Thus, a low efficiency score means a high agreement of correction outcomes across the screens. We computed gene-wise normalised Shannon entropy across screens when considering the three possible correction outcomes outputted by the *ccr*.*GWclean* function.

We considered only the genes in common across the screens, leading to 8473 genes, and removed those found at the conjunction of two segments, totalling to 7923 genes. Then, we computed the gene-wise normalised Shannon entropy *H*_*n*_, following the formula (1):

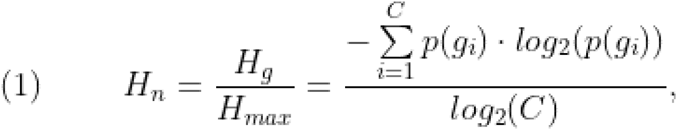

where *H*_*g*_ is the Shannon entropy of a gene *g*, computed as the sum over the variable’s probability values, *g*_*i*_ ∈ {-1, 0, 1} and *C* is the total number of possible correction outcomes (i.e. 3 in this case), divided by the maximum expected entropy (*log*_*2*_ of *C*).

#### Recall of genes in six predefined gene-sets following CRISPRcleanR correction

We assessed the effect of CRISPRcleanR correction on six predefined sets of genes, namely Hart2017 essential, Hart2014 nonessential, independent sets of essential genes derived from the MSigDB, copy number amplified genes from GDSC dataset, copy number amplified genes from CCLE dataset, and copy number amplified genes from GDSC that were not expressed (FPKM < 0.1 in HT-29).

For each screen, we computed the essentiality threshold at 5% false discovery rate (FDR) for the pre- and post-corrected screens, using the Hart2017 essential and Hart2014 nonessential as source of reference essential (EG) and nonessential genes (NEG), respectively. In particular, we ranked the gene-level depletion logFCs of the EG and NEG in increasing order. For each rank position *i*, we calculated a set of predicted fitness genes (PFG) as follows:

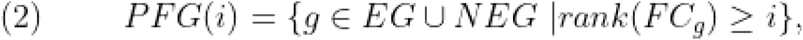

where *rank*(FC_g_) is the corresponding rank position of gene *g* in the reference gene set based on its depletion logFC. The ranked list is then used to calculate positive predictive values (PPV) for the *i*^*th*^ rank position as follows:

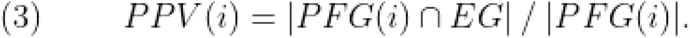

We determined the highest threshold of depletion logFC (logFC*) in rank position *i\** such that PPV(*i*^*\**^) ≥ 0.95, which is equivalent to an FDR of 5%. We considered all genes with a logFC < logFC* as essential for the viability of HT-29. For each gene-set *S*, we computed the recall by dividing the size of genes *g*_*s*_ below the 5% FDR threshold (logFC*) by the total size of *g*_*s*_ included in the screen, according to (4):

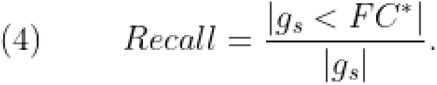

## Availability

CRISPRcleanR^WebApp^ is available at: https://crisprcleanr-webapp.fht.org (reviewers’ access credentials: username =, password = 1111 and username =, password = 1111). The latest version of the original CRISPRcleanR package can be found at the following GitHub repository: https://github.com/francescojm/CRISPRcleanR. The raw read count files obtained from six screens performed on the HT-29 cell line using different genome-wide CRISPR-Cas9 libraries are available as external data in the CRISPRcleanR package: https://github.com/francescojm/CRISPRcleanR/tree/master/inst/extdata.

## Author Contributions

AV conceived the study, designed and performed the benchmark analyses, tested the web application, wrote and revised the manuscript. RRDL designed, developed and tested the web application, integrated all the necessary libraries and technologies with the IT infrastructure, wrote and revised the manuscript. RRDL is also responsible for the maintenance and future releases of the web application. PC customised the core package to facilitate its integration in the web application and added new functionalities, tested the web application, and revised the manuscript. UP contributed to the design of the benchmark analyses, tested the web application, and revised the manuscript. EK assembled a CRISPRcleanR compatible annotation for the GeCKOv2 library and revised the manuscript. LM, CF, KHK and DSA set up the IT infrastructure and technologies required for hosting the web application at Human Technopole. FI conceived the study, designed the original algorithms and developed the R-based version of CRISPRcleanR, contributed to the design of the benchmark analyses, wrote and revised the manuscript, and supervised the study.

## Competing interests

FI receives funding from Open Targets, a public-private initiative involving academia and industry and performs consultancy for the joint CRUK-AstraZeneca Functional Genomics Centre. All other authors declare that they have no competing interests.

### Abbreviations

API: application programming interface
AUPRC: Area Under Precision-Recall Curve
AUROC: Area Under Receiver Operating Characteristic
CCLE: Cancer Cell Line Encyclopaedia
CN: copy number
CRISPR: Clustered Regularly Interspaced Short Palindromic Repeats
DSB: double strand breaks
EG: essential genes
FDR: false discovery rate
GDSC: Genomics of Drug Sensitivity in Cancer dataset
logFC: log fold-change
MSigDB: molecular signature database
NEG: nonessential genes
QC: quality control
sgRNA: single-guide RNA
PPV: positive predicted value
SPA: Single Page Application

## References

Aguirre, A.J., Meyers, R.M., Weir, B.A., Vazquez, F., Zhang, C.-Z., Ben-David, U., Cook, A., Ha, G., Harrington, W.F., Doshi, M.B., et al. (2016). Genomic Copy Number Dictates a Gene-Independent Cell Response to CRISPR/Cas9 Targeting. Cancer Discov. 6, 914–929..

Allen, F., Behan, F., Khodak, A., Iorio, F., Yusa, K., and Garnett, M. (2019). JACKS: joint analysis of CRISPR/Cas9 knockout screens. Genome Res. 29. https://doi.org/10.1101/gr.238923.118.

Barrangou, R., Birmingham, A., Wiemann, S., Beijersbergen, R.L., Hornung, V., and Smith, A. van B. (2015). Advances in CRISPR-Cas9 genome engineering: lessons learned from RNA interference. Nucleic Acids Res. 43, 3407–3419..

Barretina, J., Caponigro, G., Stransky, N., Venkatesan, K., Margolin, A.A., Kim, S., Wilson, C.J., Lehár, J., Kryukov, G.V., Sonkin, D., et al. (2012). The Cancer Cell Line Encyclopedia enables predictive modelling of anticancer drug sensitivity. Nature 483, 603–607..

Behan, F.M., Iorio, F., Picco, G., Gonçalves, E., Beaver, C.M., Migliardi, G., Santos, R., Rao, Y., Sassi, F., Pinnelli, M., et al. (2019). Prioritization of cancer therapeutic targets using CRISPR-Cas9 screens. Nature 568, 511–516..

Chai, A.W.Y., Yee, P.S., Price, S., Yee, S.M., Lee, H.M., Tiong, V.K., Gonçalves, E., Behan, F.M., Bateson, J., Gilbert, J., et al. (2020). Genome-wide CRISPR screens of oral squamous cell carcinoma reveal fitness genes in the Hippo pathway. Elife 9. https://doi.org/10.7554/eLife.57761.

Cong, L., Ran, F.A., Cox, D., Lin, S., Barretto, R., Habib, N., Hsu, P.D., Wu, X., Jiang, W., Marraffini, L.A., et al. (2013). Multiplex genome engineering using CRISPR/Cas systems. Science 339, 819–823..

Davis, A.J., and Chen, D.J. (2013). DNA double strand break repair via non-homologous end-joining. Transl. Cancer Res. 2, 130–143..

Dede, M., McLaughlin, M., Kim, E., and Hart, T. (2020). Multiplex enCas12a screens detect functional buffering among paralogs otherwise masked in monogenic Cas9 knockout screens. Genome Biol. 21, 262..

Dempster, J., Behan, F.M., Green, T., Najgebauer, H., Krill-Burger, J., Allen, F., and Others (2019a). Agreement between two large pan-cancer genome-scale CRISPR knock-out datasets. Nat. Commun. 10, 5817..

Dempster, J.M., Rossen, J., Kazachkova, M., Pan, J., Kugener, G., Root, D.E., and Tsherniak, A. (2019b). Extracting Biological Insights from the Project Achilles Genome-Scale CRISPR Screens in Cancer Cell Lines.

Dempster, J.M., Boyle, I., Vazquez, F., Root, D., Boehm, J.S., Hahn, W.C., Tsherniak, A., and McFarland, J.M. (2021). Chronos: a CRISPR cell population dynamics model.

Doench, J.G. (2018). Am I ready for CRISPR? A user’s guide to genetic screens. Nat. Rev. Genet. 19, 67–80..

Doench, J.G., Fusi, N., Sullender, M., Hegde, M., Vaimberg, E.W., and Donovan, K.F. (2016). Optimized sgRNA design to maximize activity and minimize off-target effects of CRISPR-Cas9. Nat. Biotechnol. 34. https://doi.org/10.1038/nbt.3437.

Evers, B., Jastrzebski, K., Heijmans, J.P.M., Grernrum, W., Beijersbergen, R.L., and Bernards, R. (2016). CRISPR knockout screening outperforms shRNA and CRISPRi in identifying essential genes. Nat. Biotechnol. 34, 631–633..

Gomez, V., and Hergovich, A. (2016). Chapter 14 - Cell-Cycle Control and DNA-Damage Signaling in Mammals. In Genome Stability, I. Kovalchuk, and O. Kovalchuk, eds. (Boston: Academic Press), pp. 227–242.

Gonçalves, E., Behan, F.M., Louzada, S., Arnol, D., Stronach, E.A., Yang, F., Yusa, K., Stegle, O., Iorio, F., and Garnett, M.J. (2019). Structural rearrangements generate cell-specific, gene-independent CRISPR-Cas9 loss of fitness effects. Genome Biol. 20, 27..

Gonçalves, E., Segura-Cabrera, A., Pacini, C., Picco, G., Behan, F.M., Jaaks, P., Coker, E.A., van der Meer, D., Barthorpe, A., Lightfoot, H., et al. (2020). Drug mechanism-of-action discovery through the integration of pharmacological and CRISPR screens. Mol. Syst. Biol. 16, e9405..

Gonçalves, E., Thomas, M., Behan, F.M., Picco, G., Pacini, C., Allen, F., Vinceti, A., Sharma, M., Jackson, D.A., Price, S., et al. (2021). Minimal genome-wide human CRISPR-Cas9 library. Genome Biol. 22, 40..

Haeussler, M., Schönig, K., Eckert, H., Eschstruth, A., Mianné, J., Renaud, J.-B., Schneider-Maunoury, S., Shkumatava, A., Teboul, L., Kent, J., et al. (2016). Evaluation of off-target and on-target scoring algorithms and integration into the guide RNA selection tool CRISPOR. Genome Biol. 17, 148..

Hart, T., and Moffat, J. (2016). BAGEL: a computational framework for identifying essential genes from pooled library screens. BMC Bioinformatics 17. https://doi.org/10.1186/s12859-016-1015-8.

Hart, T., Brown, K.R., Sircoulomb, F., Rottapel, R., and Moffat, J. (2014). Measuring error rates in genomic perturbation screens: gold standards for human functional genomics. Mol. Syst. Biol. 10, 733..

Hart, T., Chandrashekhar, M., Aregger, M., Steinhart, Z., Brown, K.R., and MacLeod, G. (2015). High-resolution CRISPR screens reveal fitness genes and genotype-specific cancer liabilities. Cell 163. https://doi.org/10.1016/j.cell.2015.11.015.

Hart, T., Tong, A.H.Y., Chan, K., Van Leeuwen, J., Seetharaman, A., Aregger, M., Chandrashekhar, M., Hustedt, N., Seth, S., Noonan, A., et al. (2017). Evaluation and Design of Genome-Wide CRISPR/SpCas9 Knockout Screens. G3 7, 2719–2727..

Hsu, P.D., Lander, E.S., and Zhang, F. (2014). Development and applications of CRISPR-Cas9 for genome engineering. Cell 157, 1262–1278..

Imkeller, K., Ambrosi, G., Boutros, M., and Huber, W. (2020). gscreend: modelling asymmetric count ratios in CRISPR screens to decrease experiment size and improve phenotype detection. Genome Biol. 21, 53..

Iorio, F., Knijnenburg, T.A., Vis, D.J., Bignell, G.R., Menden, M.P., Schubert, M., Aben, N., Gonçalves, E., Barthorpe, S., Lightfoot, H., et al. (2016). A Landscape of Pharmacogenomic Interactions in Cancer. Cell 166, 740–754..

Iorio, F., Behan, F.M., Gonçalves, E., Bhosle, S.G., Chen, E., Shepherd, R., Beaver, C., Ansari, R., Pooley, R., Wilkinson, P., et al. (2018). Unsupervised correction of gene-independent cell responses to CRISPR-Cas9 targeting. BMC Genomics 19, 604..

Jinek, M., Chylinski, K., Fonfara, I., Hauer, M., Doudna, J.A., and Charpentier, E. (2012). A programmable dual-RNA-guided DNA endonuclease in adaptive bacterial immunity. Science 337, 816–821..

Kim, E., and Hart, T. (2021). Improved analysis of CRISPR fitness screens and reduced off-target effects with the BAGEL2 gene essentiality classifier. Genome Med. 13, 2..

Korkmaz, G., Lopes, R., Ugalde, A.P., Nevedomskaya, E., Han, R., Myacheva, K., Zwart, W., Elkon, R., and Agami, R. (2016). Functional genetic screens for enhancer elements in the human genome using CRISPR-Cas9. Nat. Biotechnol. 34, 192–198..

Kurata, M., Yamamoto, K., Moriarity, B.S., Kitagawa, M., and Largaespada, D.A. (2018). CRISPR/Cas9 library screening for drug target discovery. J. Hum. Genet. 63, 179–186..

Lawrence, M., Huber, W., Pagès, H., Aboyoun, P., Carlson, M., Gentleman, R., Morgan, M.T., and Carey, V.J. (2013). Software for computing and annotating genomic ranges. PLoS Comput. Biol. 9, e1003118..

Li, W., Xu, H., Xiao, T., Cong, L., Love, M.I., Zhang, F., Irizarry, R.A., Liu, J.S., Brown, M., and Liu, X.S. (2014). MAGeCK enables robust identification of essential genes from genome-scale CRISPR/Cas9 knockout screens. Genome Biol. 15, 554..

Li, W., Köster, J., Xu, H., Chen, C.-H., Xiao, T., Liu, J.S., Brown, M., and Liu, X.S. (2015). Quality control, modeling, and visualization of CRISPR screens with MAGeCK-VISPR. Genome Biol. 16, 281..

Liao, Y., Smyth, G.K., and Shi, W. (2019). The R package Rsubread is easier, faster, cheaper and better for alignment and quantification of RNA sequencing reads. Nucleic Acids Res. 47, e47..

Lord, C.J., Quinn, N., and Ryan, C.J. (2020). Integrative analysis of large-scale loss-of-function screens identifies robust cancer-associated genetic interactions. Elife 9, e58925..

Mali, P., Yang, L., Esvelt, K.M., Aach, J., Guell, M., DiCarlo, J.E., Norville, J.E., and Church, G.M. (2013). RNA-guided human genome engineering via Cas9. Science 339, 823–826..

Martinez-Lage, M., Puig-Serra, P., Menendez, P., Torres-Ruiz, R., and Rodriguez-Perales, S. (2018). CRISPR/Cas9 for Cancer Therapy: Hopes and Challenges. Biomedicines 6. https://doi.org/10.3390/biomedicines6040105.

Maza, E., Frasse, P., Senin, P., Bouzayen, M., and Zouine, M. (2013). Comparison of normalization methods for differential gene expression analysis in RNA-Seq experiments: A matter of relative size of studied transcriptomes. Commun. Integr. Biol. 6, e25849..

Mermel, C.H., Schumacher, S.E., Hill, B., Meyerson, M.L., Beroukhim, R., and Getz, G. (2011). GISTIC2.0 facilitates sensitive and confident localization of the targets of focal somatic copy-number alteration in human cancers. Genome Biol. 12, R41..

Meyers, R.M., Bryan, J.G., McFarland, J.M., Weir, B.A., Sizemore, A.E., Xu, H., Dharia, N.V., Montgomery, P.G., Cowley, G.S., Pantel, S., et al. (2017). Computational correction of copy number effect improves specificity of CRISPR-Cas9 essentiality screens in cancer cells. Nat. Genet. 49, 1779–1784..

Munoz, D.M., Cassiani, P.J., Li, L., Billy, E., Korn, J.M., Jones, M.D., Golji, J., Ruddy, D.A., Yu, K., McAllister, G., et al. (2016). CRISPR Screens Provide a Comprehensive Assessment of Cancer Vulnerabilities but Generate False-Positive Hits for Highly Amplified Genomic Regions. Cancer Discov. 6, 900–913..

Olshen, A.B., Venkatraman, E.S., Lucito, R., and Wigler, M. (2004). Circular binary segmentation for the analysis of array-based DNA copy number data. Biostatistics 5, 557–572..

Ong, S.H., Li, Y., Koike-Yusa, H., and Yusa, K. (2017). Optimised metrics for CRISPR-KO screens with second-generation gRNA libraries. Sci. Rep. 7, 7384..

Pacini, C., Dempster, J.M., Boyle, I., Gonçalves, E., Najgebauer, H., Karakoc, E., van der Meer, D., Barthorpe, A., Lightfoot, H., Jaaks, P., et al. (2021). Integrated cross-study datasets of genetic dependencies in cancer. Nat. Commun. 12, 1661..

Panier, S., and Durocher, D. (2013). Push back to respond better: regulatory inhibition of the DNA double-strand break response. Nat. Rev. Mol. Cell Biol. 14, 661–672..

Park, R.J., Wang, T., Koundakjian, D., Hultquist, J.F., Lamothe-Molina, P., Monel, B., Schumann, K., Yu, H., Krupzcak, K.M., Garcia-Beltran, W., et al. (2017). A genome-wide CRISPR screen identifies a restricted set of HIV host dependency factors. Nat. Genet. 49, 193–203..

Picco, G., Chen, E.D., Alonso, L.G., Behan, F.M., Gonçalves, E., Bignell, G., Matchan, A., Fu, B., Banerjee, R., Anderson, E., et al. (2019). Functional linkage of gene fusions to cancer cell fitness assessed by pharmacological and CRISPR-Cas9 screening. Nat. Commun. 10, 2198..

Rajagopal, N., Srinivasan, S., Kooshesh, K., Guo, Y., Edwards, M.D., Banerjee, B., Syed, T., Emons, B.J.M., Gifford, D.K., and Sherwood, R.I. (2016). High-throughput mapping of regulatory DNA. Nat. Biotechnol. 34, 167–174..

Rouet, P., Smih, F., and Jasin, M. (1994). Introduction of double-strand breaks into the genome of mouse cells by expression of a rare-cutting endonuclease. Mol. Cell. Biol. 14, 8096–8106..

Sanjana, N.E., Shalem, O., and Zhang, F. (2014). Improved vectors and genome-wide libraries for CRISPR screening. Nat. Methods 11. https://doi.org/10.1038/nmeth.3047.

Sanson, K.R., Hanna, R.E., Hegde, M., Donovan, K.F., Strand, C., and Sullender, M.E. (2018). Optimized libraries for CRISPR-Cas9 genetic screens with multiple modalities. Nat. Commun. 9. https://doi.org/10.1038/s41467-018-07901-8.

Shalem, O., Sanjana, N.E., Hartenian, E., Shi, X., Scott, D.A., and Mikkelson, T. (2014). Genome-scale CRISPR-Cas9 knockout screening in human cells. Science 343. https://doi.org/10.1126/science.1247005.

Shalem, O., Sanjana, N.E., and Zhang, F. (2015). High-throughput functional genomics using CRISPR-Cas9. Nat. Rev. Genet. 16, 299–311..

Sharma, S., Dincer, C., Weidemüller, P., Wright, G.J., and Petsalaki, E. (2020). CEN-tools: an integrative platform to identify the contexts of essential genes. Mol. Syst. Biol. 16, e9698.

Shen, M.W., Arbab, M., Hsu, J.Y., Worstell, D., Culbertson, S.J., Krabbe, O., Cassa, C.A., Liu, D.R., Gifford, D.K., and Sherwood, R.I. (2018). Predictable and precise template-free CRISPR editing of pathogenic variants. Nature 563, 646–651..

Smith, I., Greenside, P.G., Natoli, T., Lahr, D.L., Wadden, D., Tirosh, I., Narayan, R., Root, D.E., Golub, T.R., Subramanian, A., et al. (2017). Evaluation of RNAi and CRISPR technologies by large-scale gene expression profiling in the Connectivity Map. PLoS Biol. 15, e2003213..

Subramanian, A., Tamayo, P., Mootha, V.K., Mukherjee, S., Ebert, B.L., Gillette, M.A., Paulovich, A., Pomeroy, S.L., Golub, T.R., Lander, E.S., et al. (2005). Gene set enrichment analysis: a knowledge-based approach for interpreting genome-wide expression profiles. Proc. Natl. Acad. Sci. U. S. A. 102, 15545–15550..

Symington, L.S., and Gautier, J. (2011). Double-strand break end resection and repair pathway choice. Annu. Rev. Genet. 45, 247–271..

Tsherniak, A., Vazquez, F., Montgomery, P.G., Weir, B.A., Kryukov, G., Cowley, G.S., Gill, S., Harrington, W.F., Pantel, S., Krill-Burger, J.M., et al. (2017). Defining a Cancer Dependency Map. Cell 170, 564–576.e16..

Tzelepis, K., Koike-Yusa, H., Braekeleer, E., Li, Y., Metzakopian, E., and Dovey, O.M. (2016). A CRISPR dropout screen identifies genetic vulnerabilities and therapeutic targets in acute myeloid leukemia. Cell Rep. 17. https://doi.org/10.1016/j.celrep.2016.09.079.

Venkatraman, E.S., and Olshen, A.B. (2007). A faster circular binary segmentation algorithm for the analysis of array CGH data. Bioinformatics 23, 657–663..

Vinceti, A., Karakoc, E., Pacini, C., Perron, U., De Lucia, R.R., Garnett, M.J., and Iorio, F. (2021). CoRe: a robustly benchmarked R package for identifying core-fitness genes in genome-wide pooled CRISPR-Cas9 screens. BMC Genomics 22, 828..

Vinceti, A., Perron, U., Trastulla, L., and Iorio, F. (2022). Reduced gene templates for supervised analysis of scale-limited CRISPR-Cas9 fitness screens. Cell Rep. 40, 111145..

Wang, T., Wei, J.J., Sabatini, D.M., and Lander, E.S. (2014). Genetic screens in human cells using the CRISPR-Cas9 system. Science 343. https://doi.org/10.1126/science.1246981.

Wang, T., Birsoy, K., Hughes, N.W., Krupczak, K.M., Post, Y., Wei, J.J., Lander, E.S., and Sabatini, D.M. (2015). Identification and characterization of essential genes in the human genome. Science 350, 1096–1101..

de Weck, A., Golji, J., Jones, M.D., Korn, J.M., Billy, E., McDonald, E.R., 3rd, Schmelzle, T., Bitter, H., and Kauffmann, A. (2018). Correction of copy number induced false positives in CRISPR screens. PLoS Comput. Biol. 14, e1006279..

Yu, C., Luo, D., Yu, J., Zhang, M., Zheng, X., Xu, G., Wang, J., Wang, H., Xu, Y., Jiang, K., et al. (2022). Genome-wide CRISPR-cas9 knockout screening identifies GRB7 as a driver for MEK inhibitor resistance in KRAS mutant colon cancer. Oncogene 41, 191–203..

Zeng, Z., Zhang, X., Jiang, C.-Q., Zhang, Y.-G., Wu, X., Li, J., Tang, S., Li, L., Gu, L.-J., Xie, X.-Y., et al. (2022). Identifying novel therapeutic targets in gastric cancer using genome-wide CRISPR-Cas9 screening. Oncogene 41, 2069–2078..

Zhou, Y., Zhu, S., Cai, C., Yuan, P., Li, C., Huang, Y., and Wei, W. (2014). High-throughput screening of a CRISPR/Cas9 library for functional genomics in human cells. Nature 509, 487–491..

