## Supplemental Information for "*CRISPRcleanR*^*WebApp*^: an interactive web application for processing genome-wide pooled CRISPR-Cas9 viability screens"

Additional documentation for

### 1 Application overall architecture

All services related to the application run on Docker (https://www.docker.com/) containers hosted by the IT infrastructure of Human Technopole. Fig. 1 shows a high level representation of the overall application.

The front-end web application runs on the client browser (e.g. Chrome, Firefox), and is served through an Nginx (https://www.nginx.com/) containerized server. Users login, registration and private data related requests are managed through Keycloak (https://www.keycloak.org/), a flexible Single Sign-On authentication+authorization service.

The backend is implemented as a microservice architecture where multiple containerized services communicate to each other through a Redis (https://redis.io/) message broker. Two servers are exposed to the client. A FastAPI (https://fastapi.tiangolo.com/) API server is used to manage the submission of new jobs and to retrieve job results data that are stored in a MongoDB (https://www.mongodb.com/) database instance. A second NodeJS (https://nodejs.org/en/) server is used to manage file exchange with the client. Files are stored on an on-premise S3 bucket (https://aws.amazon.com/s3/) compliant object storage. Jobs are processed through a dedicated Celery (https://github.com/celery/celery) tasks queue.

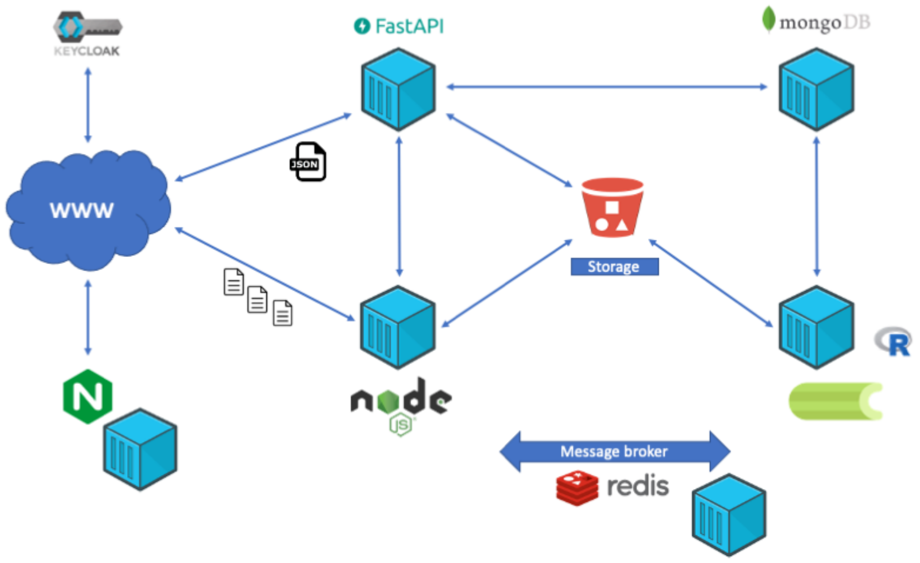


**Fig. 1 - High-level architecture of CRISPRcleanR$^{\text{WebApp}}$.**

### *1.1 Front-end*

The front-end web application is the software that runs directly on the client browser, accessible at https://crisprcleanr-webapp.fht.org/home. This front-end is designed with Vue.js, a well-known Single Page Application (SPA) framework. SPAs are entirely rendered on the client browser. Once the client hits the application URL for the first time, the Nginx server sends the entire application as a unique HTML+CSS+Javascript bundle. Client side routing, based on Javascript, is then used to render on-the-fly the HTML, depending on the current navigated page. In the meantime, dynamic data needed to fill the page (such as job results) is retrieved asynchronously from the backend, in JSON format.

The main parts of the application are constituted by two components: the multistep form and the file uploader.

The multistep form component is able to modify its appearance according to user selections and current step, and is responsible for sending job data to the backend upon a job submission. Each form step undergoes an input validation process that ensures no information is missing/ wrong during the form progression. The form evolution and validation is managed through an internal state machine implemented through the XState library (https://xstate.js.org/). The state machine describes all possible configurations of the form, keeps the current form state, and stores the input data inserted by the client. The GUI reads the machine state and renders the current form step and configuration accordingly, while communicating back to the machine user inputs via dedicated events.

The file upload component is page-independent, and it is able to manage and monitor the file upload status. Multiple files from multiple jobs can be updated in parallel. Each file upload progress is shown on the component. In order to implement such a component, uploading files related data is stored on a dedicated VueX (https://vuex.vuejs.org/) centralized storage, accessible from everywhere in the app. This allows the multi-step form to send information about each new submitted job related file to be uploaded. The upload component is constantly fed with information from the storage and triggers a new upload for each new file. Each upload is, again, managed with a state machine that monitors the upload status and provides graphical information to the upload component to show the current upload status and progress.

Keycloak JS adapter (https://github.com/keycloak/keycloak-documentation/blob/main/securing_apps/topics/oidc/javascript-adapter.adoc) has been used to implement the communication with the Keycloak server, needed to obtain JWT tokens used to authorize protected operations on the back-end, such as job upload and inspection. The obtained token is added to the HTTP header of each backend request (both API server and file server) and is validated server-side.

As far as dynamic charts are concerned, a mixed strategy which takes advantage of Vue components and D3.js (https://d3js.org/) has been used to render charts, with a particular attention to real time rerender performances.

#### *1.2 Back-end: API Server*

The API server is the entrypoint for the back-end. Communication primarily occurs through the front-end application. The API server exposes many endpoints that allow the user to:

- Download example data
- Submit new jobs
- Retrieve the jobs results list
- Retrieve single result details

Our API server is based on the python FastAPI framework. FastAPI adopts the ASGI protocol over the older WSGI one (common to frameworks such as Django and Flask), which fully exploits python coroutines to enable asynchronous Input/Output (IO) operations. Coroutines allow IO bound tasks to yield, while waiting for the IO operation to conclude, allowing other CPU bound tasks to carry on with calculations on the same process. At the IO operations completion, the task is resumed and all further operations are carried on. The better usage of computational cycles and reduction of idle times allows the ASGI protocol to efficiently handle fast-serving high-load requests with great amounts of IO tasks. In addition, the FastAPI framework allows for an easy and automatic API documentation mechanism based on swagger (https://swagger.io/) and OpenAPI schema (https://www.openapis.org/). CRISPRcleanR API server stores jobs and files information within a dedicated MongoDB instance.

#### *1.3 Back-end: Celery Tasks Queue*

Actual computation of jobs through the R package happens on a separated process within the API server, allowing us to obtain a great responsiveness from the API server, since the latter doesn't have to dedicate computational resources to elaborate jobs. The Celery tasks queue is implemented as a set of containerized workers listening for messages on a queue fed through a Redis message broker. For each new job, the API server publishes a new message to the broker, which is enqueued. An available worker reads the message with the job details and processes the job through a quite articulated task sequence, described in detail in Section 2.2.

#### *1.4 Back-end: File Server*

FastAPI file management is not flexible enough to handle large file uploads combined with upload progress monitoring and direct upload to S3. Our file server is therefore implemented as a separate entity based on NodeJS, where optimized endpoints are able to directly stream the file to S3, thanks to the BusBoy (https://github.com/mscdex/busboy) library, without the need to locally store the file on the server.

In order to make the API server aware of the files being uploaded, which is necessary for the app to decide when a job computation can be forwarded to Celery, the file server publishes a new message to the subscribed API server on a dedicated Redis message queue. The message exchange is again described in Section 2.2

#### 1.5 Authorization

CRISPRcleanR$^{WebApp}$ provides user authentication and authorization to send/retrieve data in order to maintain data private to users. The authorization mechanism follows the Authorization Code Flow with Public Client and JSON web token described within the OAuth 2.0 standard (https://oauth.net/2/grant-types/authorization-code/), and is encapsulated into the OIDC standard (https://openid.net/connect/). A Keycloak server is used to perform the authentication against an internal managed users database. An access token is issued to the client if the user authentication is completed correctly. New users can also register from the dedicated page reachable from the *Sign In or Register* section of the web app.
For each request to the back-end, both the API server or file server, the client sends the token as a bearer token, in the HTTP header, which is then validated from the server against a public certificate retrieved from the Keycloak server itself. If the token is valid, the server executes the client action (e.g. submitting a new job or files, retrieving job data).

### 2 Web app job computation

This section is devoted to providing an in-depth overview of the most important phases involved in a new job computation, from job submission to results visualization.

#### 2.1 User interaction

Fig. 2 depicts the user workflow. A user starts by filling out the webapp form with all required fields and files.

During form filling, each form step undergoes a validation to assert whether its fields contain expected data. In case of wrong or missing input, an explanatory error message is prompted below the involved input field. When a user submits the form, textual data is sent to the API server. It is worth noting that, during this phase, only file names are sent, not the actual files. A notification appears on screen with the job submission outcome: this can be either a green notification confirming the correct job submission, or a red notification with an error message, should the submission process fail for some reason (e.g. network error, server error, etc). The submitted job then appears in the application's "Results" section, along with the job status (pending, completed, error). If an error occurred during job submission, the job is immediately put in an error state, and an error chain of tasks is executed, as described in Session 2.2.

On the other hand, a “successful submission” server response contains a list of file IDs, related to the job files, that the app immediately sends to the file server as metadata, along with the actual files. A file upload component appears on screen allowing to monitor the file upload status, and giving the opportunity to cancel ongoing uploads. During this phase, uploads can again be completed successfully or go in an error state. In case of error, the job is again immediately tagged as unsuccessful and follows the error chain of tasks.

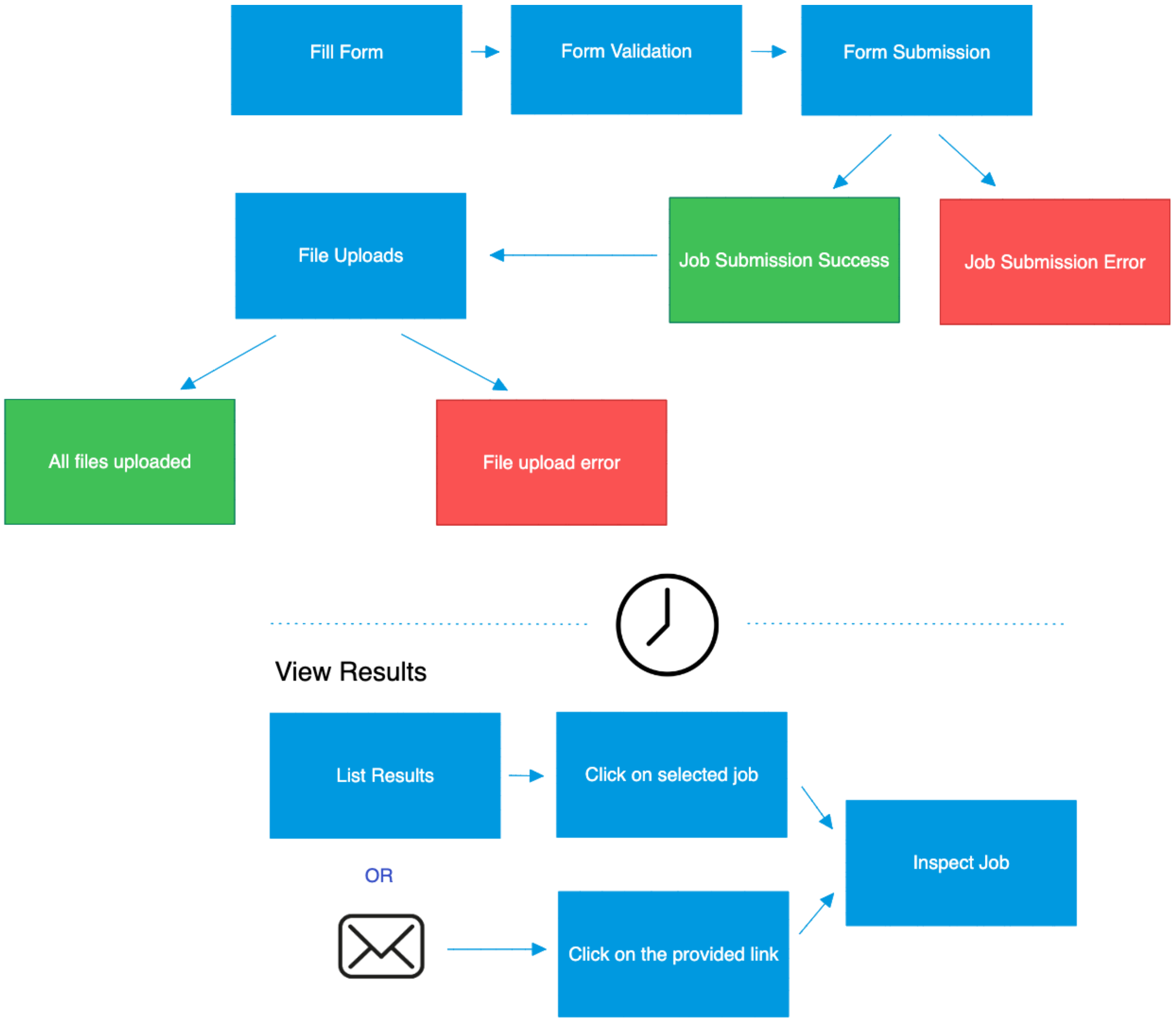


**Fig. 2 - Front-end Job submission and Results visualization workflow.**

If all file uploads have successfully finished, the API server forwards a computation request to the Celery queue, which starts the computation with the CRISPRcleanR package. Job duration depends on the back-end load, the job type (e.g. FASTQ alignment or usage of precomputed count files), the size of the files, etc. Once the computation has finished, if the user flagged his/her email address in the job submission form, s/he will receive an email notification containing the job's outcome. Success emails contain a link pointing to the detailed job results page. In case of error, this email contains a possible cause for error. This is especially useful when errors occurred during the file processing with the R package, since it can give a hint about what went wrong (file content, format, etc). For users who do not wish to receive an email, they can still manually access the job from the "Results" section.

#### *2.2 Back-end operations*

The back-end exposes a REST API server which is used to communicate with the clients for receiving and computing jobs, and returning results. Fig. 3 shows the entire flow of the back-end operations involved in a job submission.

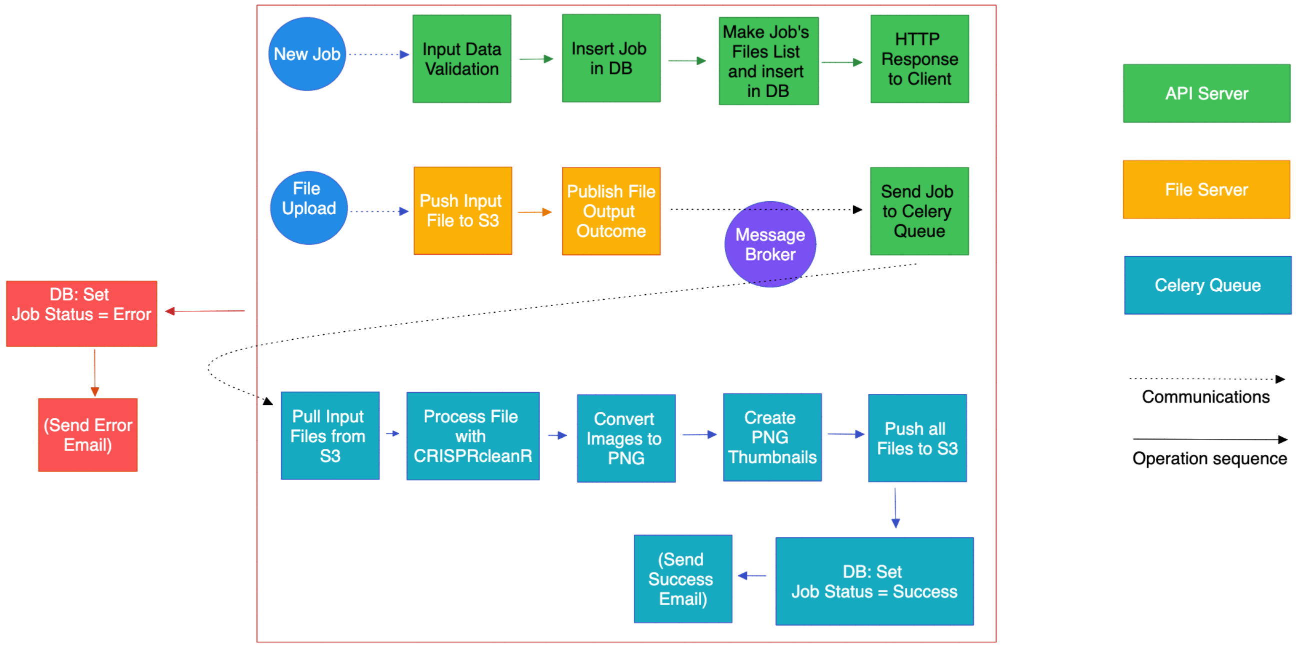


**Fig. 3 - Back-end operations for a job submission.**

Job handling starts from the API REST server, which receives a new job from the client HTTP POST request. First, form data is validated against an expected input schema in order to prevent corrupted or incorrect requests. Although the front-end already provides form validation, this has to be repeated, since the API server is an independent entity that could potentially be used by many different client applications, scripts, etc. In case of negative validation, the server immediately replies back with "code 422: Unprocessable entity", and the processing chain is put in an error state. Whenever an error occurs, an error chain of tasks is executed which performs standard operations, such as updating existing database entries with error status, and optionally sending an email to inform the user about the error. In case of positive validation, the server creates a new entry for the job in the database. The unique record ID produced by the database is then used as a unique identifier for the job. In addition, all files are inserted in a dedicated collection of the database, and their corresponding unique IDs are sent back to the user as a file list, along with the job ID. The combination between job ID and file ID is used to have a unique, job-grouped identification system for targeting files to be uploaded in the S3 bucket. The client then starts the upload of each file and sends the additional metadata about the previously described identifiers. For each upload completion or error, a message is forwarded from the file server to the API server through the message broker, which will contain useful information to understand if the upload was completed successfully for which file. When a new message is received, the API server uploads the file entries in the database with the updated status for the files. If the received file communicates an upload error, all files are put in an error state. The processing chain goes in error state and proceeds with the error chain of tasks. On the other hand, if there are still pending uploads for the current job, the API server simply updates the current file status and yields. Finally, if all jobs have been uploaded, the API server sends a message to the Celery queue asking for processing the current job.

Within the celery worker which took charge of the job, a number of subtasks are executed. First, it retrieves jobs data from the database and downloads files from the S3 bucket into a local folder. Then, the R package is called to process the job. All produced files are stored in the same local folder where input files were previously downloaded. After the successful processing, produced pdf images are duplicated in png format, low sized png thumbnails are computed out of them, and zip files are created for output data and images. Finally, output data and images are uploaded back to the S3 bucket, database entries are updated, the local folder is removed and an optional email is sent to the user. If any error occurs during these tasks, the chain is put in an error state and the error chain of tasks is executed.

At any time, the user can list all jobs submitted in the designated results section. When clicking on a particular job result, a dedicated page is loaded. For completed jobs, all resources are made available here. Output data can be downloaded. Interactive charts can be visualized starting from JSON data files requested from the API server, which were created during the R processing phase and subsequently uploaded to the S3 bucket together with the results files.
